# Discovering chromatin dysregulation induced by protein-coding perturbations at scale

**DOI:** 10.1101/2023.09.20.555752

**Authors:** Max Frenkel, Margaux L.A. Hujoel, Zachary Morris, Srivatsan Raman

**Affiliations:** Cellular and Molecular Biology Graduate Program, University of Wisconsin, Madison, Wisconsin, USA; Medical Scientist Training Program, University of Wisconsin School of Medicine and Public Health, Madison, Wisconsin, USA; Department of Biochemistry, University of Wisconsin, Madison, Wisconsin, USA; Division of Genetics, Department of Medicine, Brigham and Women’s Hospital and Harvard Medical School, Boston, MA, USA; Center for Data Sciences, Brigham and Women’s Hospital and Harvard Medical School, Boston, MA, USA; Program in Medical and Population Genetics, Broad Institute of MIT and Harvard, Cambridge, MA, USA; Department of Human Oncology, University of Wisconsin School of Medicine and Public Health, Madison, Wisconsin, USA; Department of Bacteriology, University of Wisconsin, Madison, Wisconsin, USA; Department of Chemical and Biological Engineering, University of Wisconsin, Madison, Wisconsin, USA

## Abstract

Although population-scale databases have expanded to millions of protein-coding variants, insight into variant mechanisms has not kept pace. We present PROD-ATAC, a high-throughput method for discovering the effects of protein-coding variants on chromatin. A pooled library of variants is expressed in a disease-agnostic cell line, and single-cell ATAC resolves each variant’s effect on chromatin. Using PROD-ATAC, we characterized the effects of >100 oncofusions (a class of cancer-causing chimeric proteins) and controls and revealed that pioneer activity is a common feature of fusions spanning an enormous range of fusion frequencies. Further, fusion-induced dysregulation can be context-agnostic as observed mechanisms often overlapped with cancer and cell-type specific prior knowledge. We also showed that gain-of-function pioneering is common among oncofusions. This work provides a global view of fusion-induced chromatin. We uncovered convergent mechanisms among disparate oncofusions and shared modes of dysregulation across different cancers. PROD-ATAC is generalizable to any set of protein-coding variants.

## Introduction

Advances in DNA sequencing have allowed geneticists to catalog millions of variants throughout the human genome in diverse populations, cell types, and disease states. At the same time, improvements in targeted genomic manipulation and DNA synthesis have allowed for the development of high-throughput methods for discovering variant effects. These methods achieve scale by almost universally sacrificing resolution. For instance, massively parallel reporter assays can measure between 10^3^ and 10^6^ variants per experiment by collapsing each variant’s effects onto a single reporter^1–5^. However, one putatively representative reporter may belie complexity especially for transcription factors and chromatin regulators which execute their functions differentially and widely across the genome. Similarly, assays based on selecting for cells with functional variants are rapidly scalable but are restricted to highly abstract phenotypes like cell viability^6^ or protein stability^7^ which limit our ability to define complex genomic mechanisms. On the other hand, deep phenotypic measurements like bulk RNA or ATAC sequencing can resolve global genomic complexity but are relegated to low-throughput, arrayed reverse genetics experiments. In several cases, the scale of pooled assays has been coupled to high resolution readouts like single-cell RNA and ATAC sequencing. In these cases, CRISPR guide RNAs serve as perturbations and single-cell sequencing serves as a variant-agonistic readout for resolving complex, genome-wide effects^8–12^. Only one method exists for annotating protein-coding (instead of CRISPR) perturbations by their influences on gene expression^13^. There is no comparable scalable method for understanding the mechanisms by which protein-coding variants alter epigenetic measures of cell state and regulation at scale.

We created a scalable method that is both variant and disease agnostic for mapping the effects of protein-coding (<u>PR</u>otein-c<u>OD</u>ing) variants to their chromatin perturbations using single-cell <u>ATAC</u> (PROD-ATAC). This method achieves both the scale of pooled assays and the resolution of deep phenotyping used in traditional bulk reverse genetics experiments. Here, libraries of protein-coding variants are expressed in a pooled format and a common cellular context (clonal 293T cells). The assay’s pooled nature allows users to profile hundreds to thousands of unique variants simultaneously. The common cellular context means that each variant represents a single, well-controlled reverse genetics experiment for unambiguously establishing causal mechanisms. To resolve each variant’s effect on global chromatin architecture, we modified an existing CRISPR-based single-cell ATAC sequencing method, Spear-ATAC^14^, to capture chromatin accessibility and the encoded variant from individual cells. These modifications generalize the method beyond small CRISPR guide RNAs to accommodate protein-coding libraries of arbitrary size and complexity. PROD-ATAC therefore allows controlled expression of protein-coding variants and retrieval of each variant’s impact on the chromatin landscape broadly (Fig 1a).

**Figure 1:**
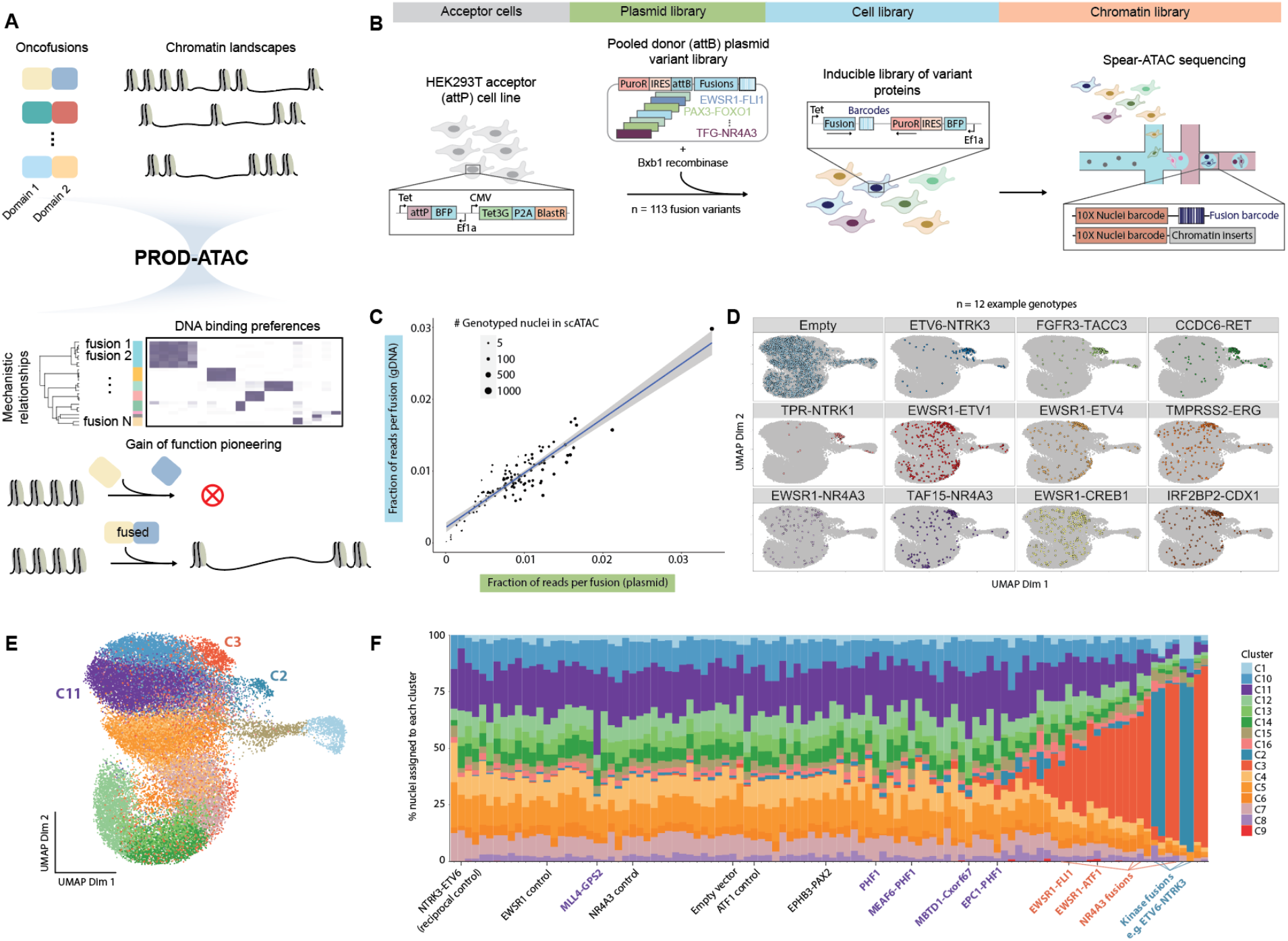
PROD-ATAC is a generalizable method for discovery of protein-coding perturbation effects at scale. (A) Overview of high-throughput mechanism discovery. Protein-coding variants alter chromatin landscapes and their mechanisms are recovered in a single pooled assay. (B) Scheme for resolving individual coding effects from pooled libraries. Bxb1 recombines pooled plasmid libraries of protein-coding variants with barcodes into HEK293T cells with a pre-integrated landing pad. Variant expression is induced and Spear-ATAC resolves chromatin accessibility and perturbation identity for individual nuclei. (C) Distribution of variants in the pooled plasmid library versus genomic DNA (gDNA) post-integration. Point size is proportional to the number of nuclei genotyped in Spear-ATAC. (D) UMAP embedding after latent semantic indexing of 35,086 genotyped nuclei. Subpanels with unique colors highlight the locations in UMAP-space for 11 example variants that shift cell-state compared to empty vector control. Clusters C2, C3, and C11 are highlighted for discriminating among variant-induced chromatin states. (E) UMAP embedding of nuclei colored based on unsupervised clustering using Seurat. (F) Fraction of cells for variant (n = 106) within each of the 16 identified clusters.

We applied PROD-ATAC to reveal chromatin dysregulation caused by over 100 potential oncogenic fusion proteins and controls. Oncofusions are chimeric proteins created by translocations that fuse the coding sequences of two unrelated proteins. Often, they contain the DNA binding domain of one protein fused to a regulatory domain of another thereby generating an aberrant protein that causes cancer^15,16^. For a few recurrent examples, these *de novo* proteins dysregulate chromatin and enhancer logic widely across the genome and in complex ways that cannot be understood with single-gene reporter assays^17–19^. At the same time, as researchers have sequenced more (and particularly rare) tumor samples, the list of fusions without mechanistic annotation has grown dramatically and is now in the tens of thousands^20^. To begin addressing this and to demonstrate PROD-ATAC’s versatility, we profiled chromatin alterations induced by a diverse set of oncofusions and controls allowing us to unambiguously test causal hypotheses.

In one experiment, we revealed mechanisms for and relationships among fusions representing more than 30 unrelated cancer subtypes. Variants profiled also ranged widely in tumor frequencies spanning rare to highly recurrent. Unlike existing studies which focus on highly recurrent fusions in narrow cancer contexts, we show that chromatin pioneering is a common feature of many oncofusions regardless of tumor frequency or subtype. Profiling rare variants also allowed us to suggest there might be heterogenous chromatin dysregulation even among fusions that cause the same cancer types. On the other hand, there were also surprising examples where seemingly unrelated fusions converged on similar modes of chromatin dysregulation. These types of non-intuitive relationships are only possible to examine with high-throughput approaches like PROD-ATAC. Importantly, we also showed that cancer and cell-type relevant information can be learned even when using 293T cells as a biosensor. Finally, we leveraged PROD-ATAC’s throughput to show that many fusions exhibit gain-of-function pioneer activities not attributable to their component parts. Our systematic approach produced a global view of the oncogenic landscape across a diverse set of biological questions and contexts that are otherwise intractable to study without scalable methods.

## Results

### A disease-agnostic method for measuring the epigenetic effects of protein-coding perturbations at scale

While several methods exist for profiling CRISPR-based perturbations with deep phenotypic readouts, these are limited technically to small guide RNAs and limited biologically to contexts where the target genes are expressed. A method for discovering genetic mechanisms at scale should be broadly applicable to any set of protein-coding variants regardless of variant size or potential mechanism, and it should evaluate each variant’s effect in a common context. Functional genomics assays for screening libraries of genetic variants often rely on lentiviral delivery; however, lentivirus production is sensitive to payload length^21^, lentivirus integrates into genomes with bias that can create context-specific effects^22^, and lentiviral template switching can shuffle barcode associations particularly for lengthy sequences^23–27^. These caveats are especially problematic for libraries of protein-coding perturbations. Protein variants can be many kilobases which limits lentiviral titers and promotes barcode shuffling. Introducing context-specific biases is problematic for single-cell sequencing readouts for which it is difficult to average over enough cells to deconvolve context-specific from variant-specific effects. We addressed each of these issues by using a system based on a site-specific recombinase (Bxb1) that is orthogonal to the mammalian genome, does not suffer from barcode shuffling, has no known cargo capacity limit, and does not require a low multiplicity of infection^28^ (Fig. 1b, Supp Fig 1a-c). Pooled libraries of variants are recombined by Bxb1 into clonal 293T acceptor cell lines that contain a single copy of a pre-integrated landing site. Cells are selected for successful recombinants each containing exactly one variant controlled by an inducible expression system.

There are several challenges in retrieving single-copy encoded genotypes when capturing epigenetic information as in single-cell ATAC sequencing (scATAC-seq). Retrieving genotypes is relatively easy when genotypes are contained in expressed transcripts and single-cell RNA sequencing is the readout of choice (as in Perturb-seq)^9,10,29^. Spear-ATAC address this in the context of pooled guide RNA (gRNA) libraries by incorporating sequences (Nextera adapters and custom primer-binding sites) to amplify the gRNA perturbation alongside each nucleus’s chromatin library^14^. We adopted Spear-ATAC to generate amplicons for associating protein-coding perturbations to cells, and we modified the delivery construct to work for arbitrarily large coding sequences instead of small gRNAs. In lieu of directly capturing the gRNA, we added two molecular barcodes to each variant’s 3’ untranslated region (UTR). One barcode is synthesized with the protein-coding variant and the other is flanked by custom sequences (as in Spear-ATAC) such that it can be specifically enriched after the chromatin library is prepared. In this way, these 3’ UTR barcodes are proxies for each nucleus’s encoded protein-coding perturbation (Methods, Supp Fig 1d). By capturing and enriching for the 3’ UTR barcode in the same amplicon as the nuclear barcode (from 10X Genomics) we can create maps between each genotype and its associated high-content epigenetic phenotype using scATAC-seq.

This method allows pooled expression of arbitrary <u>PR</u>otein-c<u>OD</u>ing variants and discovery of their global effects on chromatin accessibility with single-cell <u>ATAC</u> (PROD-ATAC). We applied PROD-ATAC to determine the mechanisms by which oncofusions dysregulate chromatin at scale. While there are clear mechanisms by which a handful of well-known oncofusions disrupt chromatin^17,18,30–33^, there is a growing list of thousands for which we have no mechanistic insight. We mined the Catalogue of Somatic Mutations in Cancer (COSMIC)^34^ and ChimerDB^20^ to generate a list of oncofusions that partially represents the diversity of this unexplored space (Table 1). These include fusions with a wide range of tumor frequencies (from private to pathognomonic), known or suspected mechanisms, and derived from many histologic cancer subtypes (Methods). We added controls to test several *a priori* hypotheses and to benchmark the assay’s validity. The final library included 113 variants with tumor frequencies ranging from 0 to thousands of instances in COSMIC and as few as 1 instance in ChimerDB.

Each variant in the library was barcoded and cloned into the plasmid donor construct amenable for downstream Spear-ATAC sequencing. The pooled variant library was recombined into the clonal acceptor 293T sensor line, recombinants were selected for, and the final cell library was established with relatively uniform variant coverage which correlated well with the initial input distribution in the plasmid library (Methods, Fig 1c). Cells were induced to express their encoded variant before nuclei were prepared. Spear-ATAC sequencing was then used to create a chromatin accessibility library for 113,645 high-quality nuclei and a simultaneous dial-out genotyping library to assign nuclei to their encoded variant perturbations (Supp Fig 1i). Rigorous genotyping criteria were applied and variants were assigned unambiguously to 35,201 nuclei (31.0% which is comparable to previously published genotyping rates^14^) representing 112 of 113 (99.1%) variants (Methods). On average, there were 314 genotyped nuclei (median 284) identified per variant and the number of genotyped nuclei per variant correlated well with the distribution of variants in both the plasmid and cell libraries (Fig 1c, Supp Fig 1g). Quality control metrics including transcription start site (TSS) enrichment scores and number of fragments in peaks were similar across all genotypes except for kinase-containing fusions which had lower TSS enrichment (Supp Fig 1j-l).

To qualitatively assess the effect of expressing oncofusions on chromatin, we examined the relative distributions of cells containing each genotype across dimension-reduced space. On all high-quality nuclei, we performed iterative latent sematic indexing with ArchR^35^ followed by UMAP dimension reduction (Fig 1d). We restricted the variants examined in all subsequent analysis to those with at least 30 high-quality nuclei (35,086 total nuclei representing 106 variants), and then compared the relative distributions of each genotype from the pooled library. First, cells containing empty vector (EV) control distributed widely and without clear structure, representing naturally occurring heterogeneity in 293T culture. Heterogeneity within cell culture at single-cell resolution (even in clonal populations) has been previously reported for K562 cells^14,36^. Next, we examined the distributions of several known causal fusions which revealed that many conspicuously shift cell state and discriminate from the EV distribution. These fusions include kinases ETV6-NTRK3, TPR-NTRK1, and FGFR3-TACC3; ETS-containing fusions such as EWSR1-ETV1 and EWSR1-ETV4; NR4A3-containing fusions; and even the rare fusion IRF2BP2-CDX1 (Fig 1d). This demonstrated that some fusions exert pronounced effects on cell state that are resolvable even with low-resolution dimension reduction.

We then sought to examine fusion effects from a global perspective. After latent semantic indexing, we performed unsupervised graph-based clustering with Seurat^37^ which yielded 16 clusters (Fig 1e). By comparing each variant’s distribution across these clusters to that of empty vector (EV) control containing cells (and to cells of all other variants), we can determine which clusters represent fusion-specific effects versus naturally occurring heterogeneity in 293T cells (Fig 1f, Supp Fig 2). EV-containing cells and many of the controls were widely distributed but were almost never members of clusters C2 or C3. On the other hand, well-known known oncogenic fusion proteins (including all ETS-family containing fusions and all NR4A3-containing fusions) were often strongly redistributed with large abundances in C3. The cluster C2 was similarly fusion-specific and the four kinase-containing fusions (ETV6-NTRK3, FGFR3-TACC3, CCDC6-RET, and TPR-NTRK1) were dominant members of this cluster. Cells expressing the reciprocal fusion NTRK3-ETV6 (which does not encode a known functional protein) and the fusion EPHB3-PAX2 (which encodes a kinase that lacks a catalytically active domain) were almost never found in C2. While not as stark as C2 and C3, the five variants with the largest abundances in cluster C11 were fusions or controls that all contained polycomb-related proteins (EPC1-PHF1, MEAF6-PHF1, PHF1, MBDTD1-Cxorf67, and MLL4-GPS2). Several clusters therefore clearly discriminated fusions with shared known mechanisms and components. This suggests that oncofusion proteins disrupt chromatin and shift cell state distribution in ways consistent with their underlying mechanisms and that this redistribution is recoverable with PROD-ATAC.

### Oncofusion proteins often dysregulation chromatin accessibility

To quantitatively assess the ability for each library member to alter chromatin we compared the chromatin landscape of each variant to that of empty vector control. First, we called peaks with MACS2^38^ in all cells grouped by genotype. Next, we created pseudobulk replicates of the genotypes with which we then performed pairwise differential testing between all 105 experimental variants and EV control (Fig 2a). Fusions exhibit a wide range of abilities to alter chromatin accessibility. For well-known causal fusions, chromatin dysregulation is a common mechanism. Pathognomonic (i.e. characteristic) fusions such as EWSR1-FLI1 that causes Ewing sarcoma resulted in several thousand differentially accessible peaks as did rare fusions that cause Ewing sarcoma (e.g. EWSR1-ETV4), all fusions containing kinases, and those containing NR4A3. The fusions EWSR1-ATF1 and EWSR1-CREB1 both of which cause clear cell sarcoma also altered the accessibility of thousands of peaks. PAX7-FOXO1 which is pathognomonic for alveolar rhabdomyosarcoma induced several hundred differentially accessible peaks. Importantly, there were almost no differentially accessible peaks for many relevant controls despite capturing a similar number of nuclei as the fusions suggesting that our false discovery rate is well calibrated (Table 1). This included identifying 0 differentially accessible peaks for the reciprocal fusions ATF1-EWSR1, NTRK3-ETV6, NR4A3-EWSR1 (which do not encode known functional proteins), and the broken kinase EPHB3-PAX2. COL1A1-PDGFB was another useful negative control; given that it is a mitogen and does not contain a DNA binding domain, our prior was that it would be unlikely to alter chromatin directly, and we observed no differential peaks. This showed a wide range of abilities for unrelated oncofusions to disrupt chromatin and that our assay was well-calibrated for defining variant effects.

**Figure 2:**
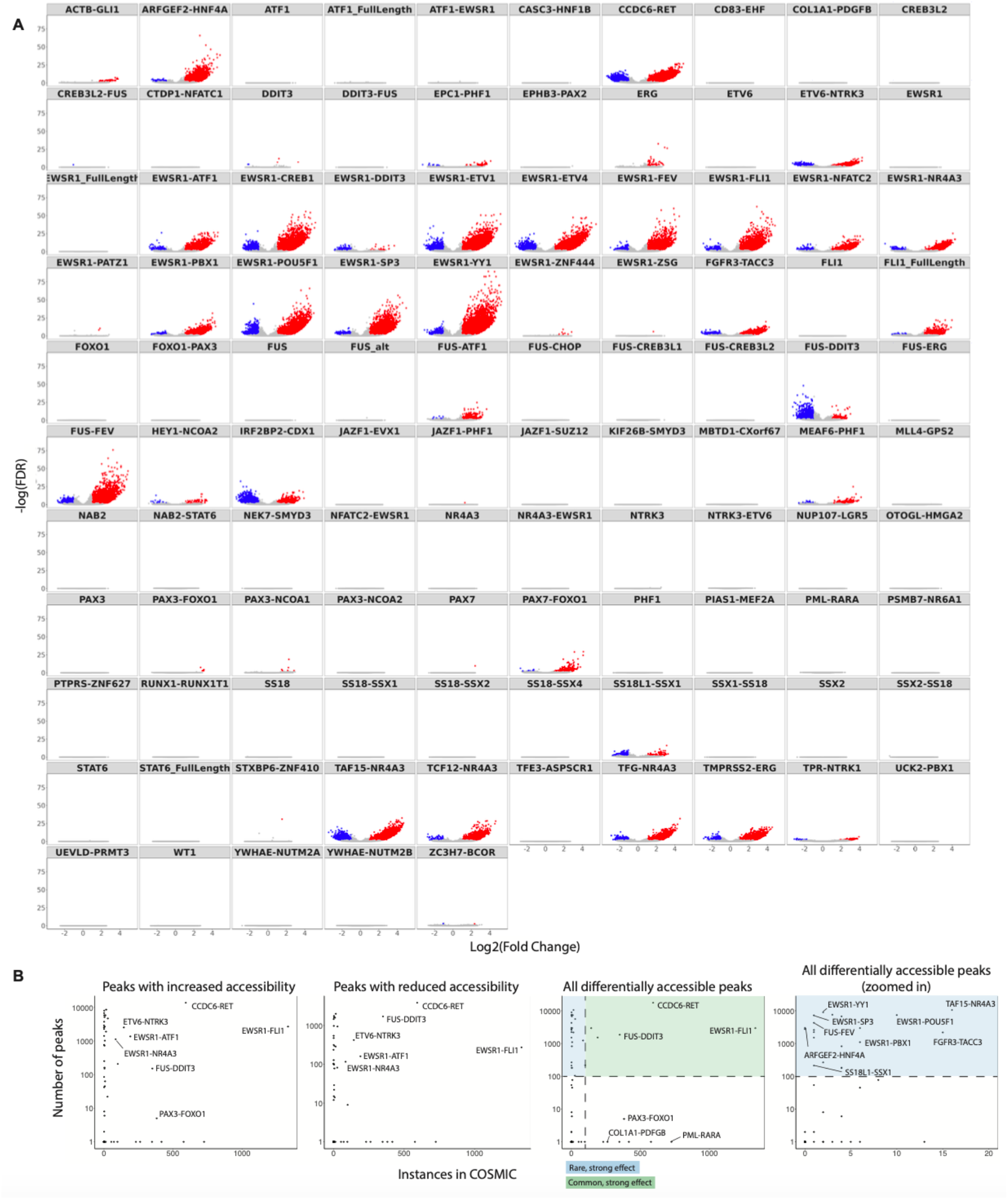
Oncofusion proteins frequently disrupt cell-state by pioneering chromatin. (A) Volcano plots for 105 variants comparing pseudobulk replicates to empty vector control. Each subplot shows -Log(False discovery rate) versus Log_2_FC for each called peak. Peaks with increased accessibility (FDR ≤ 0.1 and Log_2_FC ≥ 1) are colored red whereas those with reduced accessibility (FDR ≤ 0.1 and Log_2_FC ≤ -1) are colored blue. (B) Number of differentially accessible peaks for each fusion versus the number of instances the fusion is represented in COSMIC. Rare variants (fewer than 100 instances) with large effect size (>100 differentially accessible peaks compared to empty vector control) are highlighted blue whereas common variants with large effect size are highlighted green.

A few false negatives stood out. This included PAX3-FOXO1 which only differentially regulated 5 peaks despite being known to induce accessibility changes. When we repeated the assay with a smaller library and therefore more PAX3-FOXO1 expressing nuclei (n = 243 vs 107 nuclei, 127% increase), we saw a significant increase in the number of differentially accessible peaks (224 increased and 25 reduced peaks; Fig 2a, Supp Fig 3a). We also found that 293T natively has relatively high accessibility at known PAX3-FOXO1 target sites^39^ (Supp Fig 3b). Both low nuclei counts and natively accessible target chromatin limited our power for resolving PAX3-FOXO1’s effects (see also Fig 6 and down-sampling section). Interestingly, while most common synovial sarcoma fusions had no effect on chromatin accessibility (including SS18-SSX1, SS18-SSX2, SS18-SSX4) the rare synovial sarcoma fusion SS18L1-SSX1 *did* pioneer chromatin. We interpret this to mean that 293T cells are likely a poor model for common fusions that cause synovial sarcoma. However, there may also be heterogeneity and rare synovial sarcoma fusions may exert mechanistically distinct forms of cancer compared to their common counterparts. This is particularly interesting given that SS18L1-SSX1 and SS18-SSX1 share the same SSX1 domain and overall ∼60% of their primary amino acid sequence. Taken together, we show that chromatin dysregulation is a common feature of rare and common fusions and that our experiment is largely well-powered for discriminating variant effects.

In most cases, the preponderance of differentially accessible peaks was in the direction of increased accessibility upon expression of the fusion. This is consistent with known biases of many pioneer factors. There are two notable exceptions: FUS-DDIT3 which causes myxoid liposarcoma (1,708 or 91.9% of differential peaks were reduced in accessibility) and the rare fusion IRF2BP2-CDX1 (1,463 or 65.8% of differential peaks were reduced in accessibility). Importantly, the reciprocal fusion DDIT3-FUS and the FUS only domain control had zero differentially accessible peaks while the DDIT3 domain only control had just 5 differentially accessible peaks (despite capturing more than 500 nuclei for each genotype) again suggesting the method is well-calibrated. Chromatin dysregulation is therefore a common mechanism of oncofusions and (with interesting exceptions) there is bias towards increasing chromatin accessibility.

While PROD-ATAC revealed well-known oncofusions as chromatin regulators, it also identified effects of rare variants that would otherwise be difficult to characterize from patient samples. We plotted the number of differentially accessible peaks for each fusion against each fusion’s frequency in the COSMIC database (Fig 2b). Several rare fusions exerted dramatic changes both in increasing and decreasing chromatin accessibility at hundreds to thousands of loci. These included fusions that involve EWSR1 being fused to PBX1, POU5F1, SP3, or YY1 as well as the rare fusions IRF2BP2-CDX1, SS18L1-SSX1, ARFGEF2-HNF4A, and ACTB-GLI1. While highly recurrent fusions would be expected to dysregulate cell state as their causality is corroborated by their tumor frequency (Fig 2b left panel), rare or even patient-specific fusions appear equally capable of altering chromatin accessibility (Fig 2b right panel). Indeed, 20 of the total 73 (27.4%) non-control fusions that induced differential accessibility at more than 100 loci had fewer than 20 instances in the COSMIC database. Yet, 15 of those 20 (75%) fusions had more than 1,000 differentially accessible peaks. The fusion EWSR1-YY1 regulated the third largest number of loci (8,925 differentially accessible peaks) despite having only two instances in the COSMIC database. In summary, both rare and highly recurrent fusions can be equally capable of altering chromatin and that in most cases both are biased towards increasing accessibility.

We then sought to characterize higher-order relationships (rather than just fusion versus control) among all variants based on how they alter chromatin. We compared all 106 variants across 37,701 marker peaks and hierarchically clustered both the variants and the marker peaks (Fig 3a). This revealed clear structural relationships among the variants. Clades largely discriminated based on peaks with high z-scores (increased accessibility relative to the mean) consistent with variant bias towards inducing accessibility. Most of the control variants had few distinguishing peaks or clusters. On the other hand, all NR4A3-containing fusions, all kinases-containing fusions, all ETS-containing fusions, and all CREB family-containing fusions clearly separated into their requisite groups. These and other clades were largely separable based on the known mechanisms of the fusion components (Fig 3b) rather than the specific cancer subtype(s) they cause. For instance, all 4 kinase containing fusions strongly clustered together despite driving a wide range of cancer types and despite having a wide range of frequencies in tumor samples (e.g. CCDC6-RET, ETV6-NTRK3, and FGFR3-TACC3 are all present in > 0.1% of AACR GENIE cases whereas TPR-NTRK1 is present in only 0.04% of such cases^40^). To determine how robust these relationships are to changes in the distance metric chosen, we calculated all pairwise distances with four metrics and took the variance of those estimators to be a measure of confidence in the grouping. Indeed, the ETS, NR4A3, and CREB clades all had significantly lower variances than all other pairwise measurements (Supp Fig 4). The distance between CCDC6-RET and FGFR3-TACC3 was also robust to change in the metric used; however, there was high variance for comparisons between these and either TPR-NTRK1 or ETV6-NTRK3 (the other two kinases). Given that only 48 and 84 nuclei were captured for TPR-NTRK1 and ETV6-NTRK3, respectively (Table 1), the uncertainty in their clustering likely reflects incomplete saturation of their effects, which can be overcome with future large-scale studies designed to examine subgroup heterogeneity.

**Figure 3:**
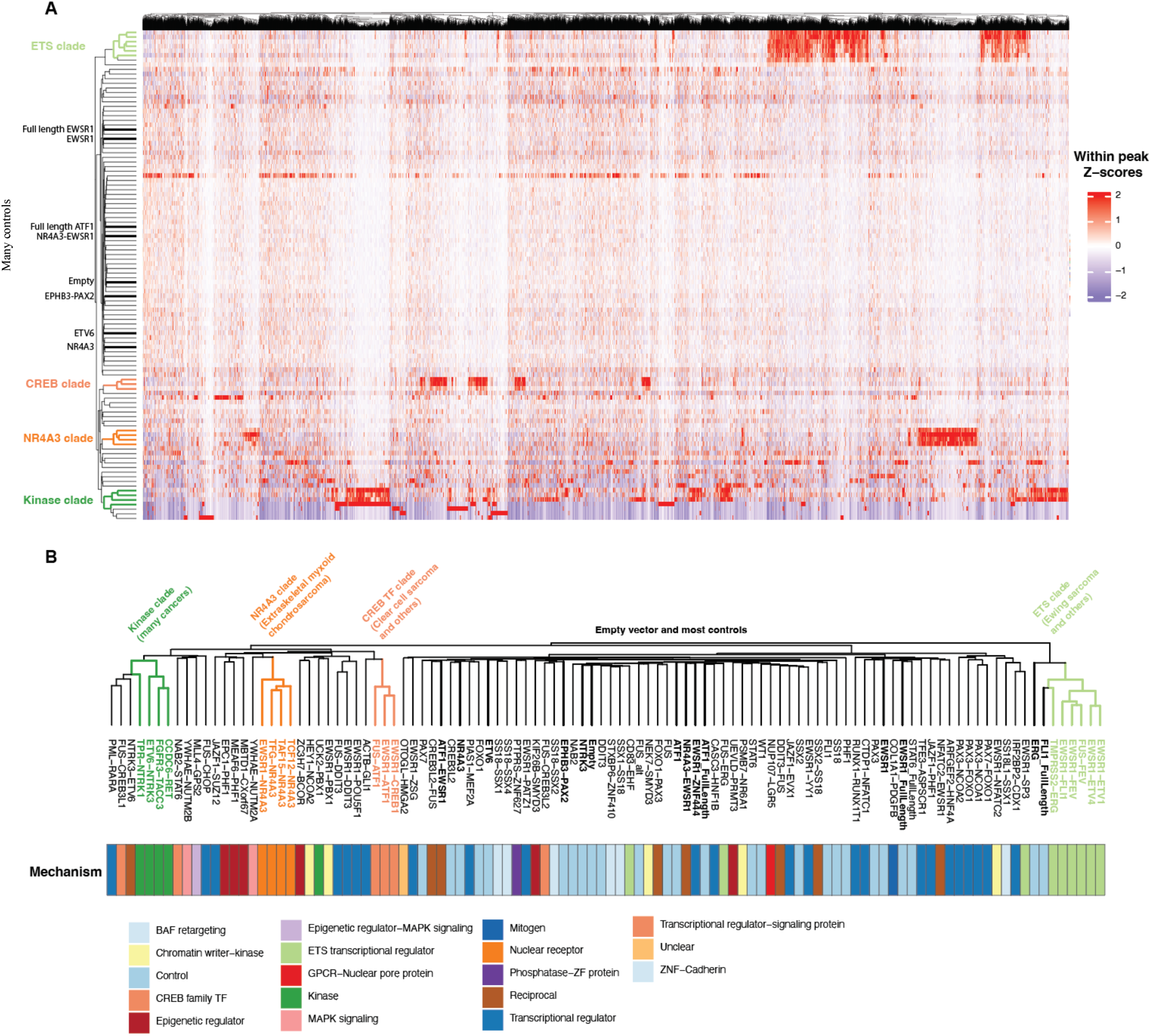
Oncofusion-expressing cells cluster based on shared mechanisms of chromatin dysregulation. (A) Heatmap of z-score normalized accessibility at 37,701 peaks that are differentially accessible (FDR ≤ 0.1 and Log_2_FC ≥1) in at least one of the variants. Columns of peaks and rows of variants are both hierarchically clustered. Clear clades that discriminated based on mechanism are colored whereas some controls are highlighted in black. (B) Hierarchical clustering of 106 variants based on z-score normalized peak accessibility by Pearson correlation (rows in panel A). Metadata is colored by known mechanism(s) of the oncofusions or their respective components. Clusters of shared mechanisms are highlighted in color in the dendrogram corresponding to those highlighted in panel A, and some relevant controls are highlighted in black.

Taken together, PROD-ATAC has revealed structure within this large library of protein variants that is consistent with known mechanisms but also revealed large scale genomic changes even for rare variants that haven’t previously been studied.

### Mechanisms of oncofusion proteins are often context-agnostic

Assaying all 106 variants in the same genetic and cellular context is a useful approach for defining causality and making well-controlled comparisons. However, it is limited in its ability to ascertain context and cell-type specific effects. Nonetheless, we reasoned that many causal oncofusions would recapitulate similar chromatin remodeling in 293T cells as they would in their respective cells of origin because they are often the sole oncogenic drivers of their cancers^15,41,42^. First, we compared pseudobulk chromatin accessibility data from our experiment to context-specific data of the well-characterized fusion EWSR1-FLI1 in Ewing sarcoma. A high-confidence set of loci bound by EWSR1-FLI1 across 15 Ewing sarcoma cancer cell lines was previously published^33^. We examined the normalized ATAC signal at these 1,879 EWSR1-FLI1 bound sites after combining all nuclei for each variant in our library (Fig 4a). Accessibility was basally low in 293T cells containing the empty vector control. Fusions with no known relation to these loci (including EWSR1-ATF1, PAX3-FOXO1, and ETV6-NTRK) had no change in accessibility, whereas there was a substantial increase in accessibility at these sites for cells expressing EWSR1-FLI1. We extended this analysis to include all ETS-family fusions most of which cause Ewing sarcoma. All ETS fusions increased accessibility at Ewing sarcoma sites and often to a greater extent than the non-fused wild type FLI1 control alone (Fig 4b). Interestingly, while EWSR1-FLI1 is the most well-known cause of Ewing sarcoma and is present in 0.23% of AACR GENIE cases (1,333 instances in COSMIC), the fusions of EWSR1 with ETV1, ETV4, and FEV all cause similarly increased accessibility at the same loci despite being less common (4, 3, and 6 instances in COSMIC and 0.13%, 0.09%, and 0.04% of AACR GENIE cases, respectively). The same analysis for PAX3-FOXO1 expressing 293T cells at loci defined in rhabdomyosarcoma cells lines^39^ and for EWSR1-ATF1 expressing 293T cells at loci defined in clear cell sarcoma cell lines^43^ revealed a similar increase in accessibility at cancer-defined loci even in 293T cells, especially when a large number of nuclei were measured (Supp Fig 3b,c). Together this shows that despite the non-native context of 293T cells, oncofusion overexpression can recapitulate chromatin alterations found in diverse cancer contexts.

**Figure 4:**
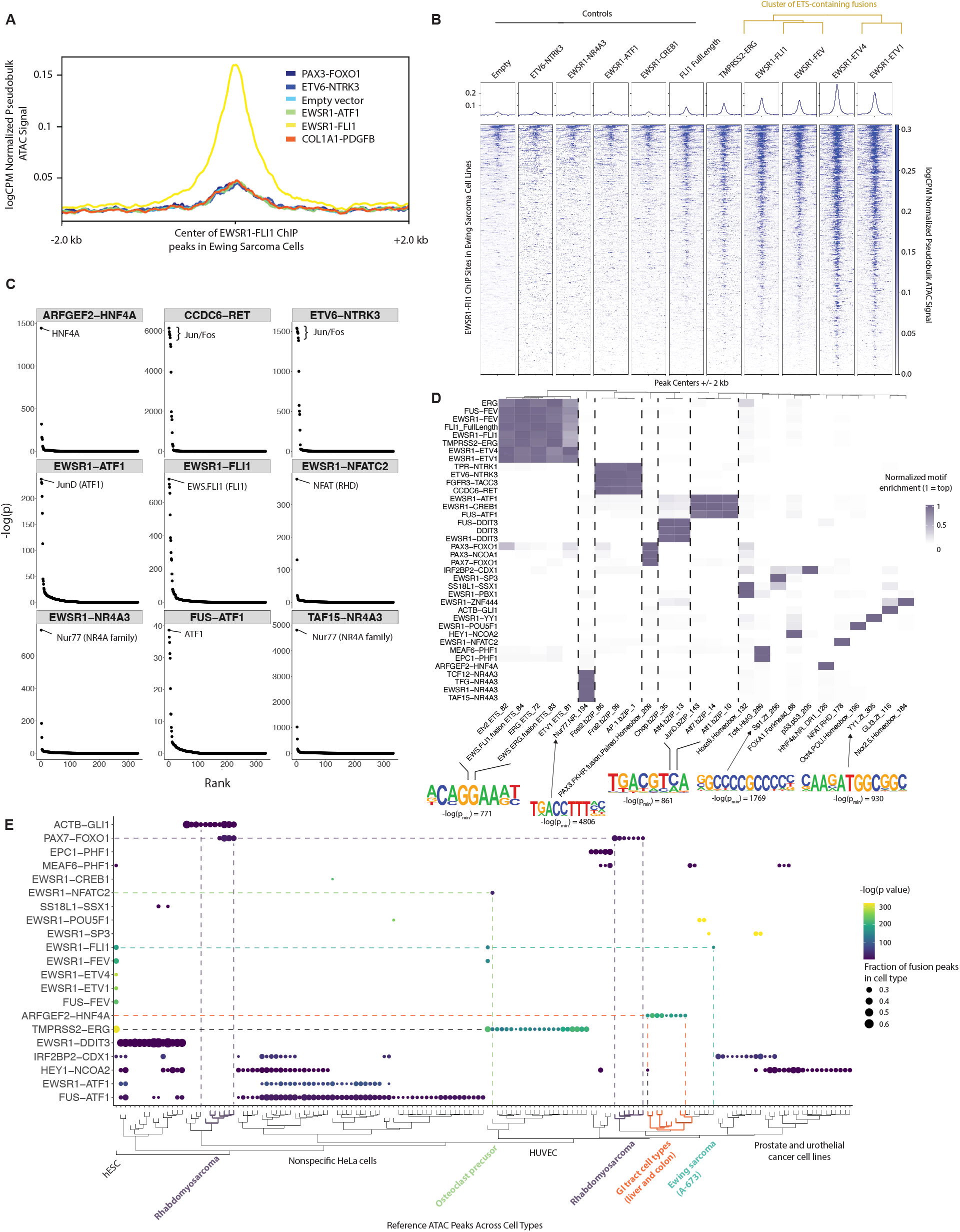
Recapitulation of known oncofusion mechanisms and discovery of novel chromatin disruption. (A) Metaplot comparing log(counts per million) (logCPM) normalized ATAC signal from pseudobulk samples of cells expressing one of 5 different fusions and empty vector (EV) control. Plot is centered at 1,879 peaks defined by EWSR1-FLI1 ChIP-Seq in Ewing sarcoma cell lines from Orth et al. 2022 (B) Metaplot and tornado plots for the same 1,879 Ewing sarcoma-specific sites across pseudobulk samples expressing one of 10 different variants or EV control. ETS-containing fusions are clustered as in Figure 3. (C) Representative DNA motif enrichment within increased differentially accessible peaks for 9 different oncofusion variants. -log(adjusted p values) plotted versus rank order for all queried motifs. (D) Clustering of motif enrichment across all fusions with enriched motifs. Within each comparison of variant to EV control, the adjusted p value of enrichment for all DNA motifs was normalized to that of the most significant DNA motif. Only variants with at least one enriched motif (p value < 0.01) and only motifs enriched in at least one pseudobulk sample are displayed. Motifs and variants were both clustered by Euclidean distance. P_min_ is the smallest adjusted p-value for a given motif. (E) Overlaps between differentially accessible peaks in pseudobulk for each fusion and significant ATAC peaks for cell types from ChIP-Atlas. Size of the dot corresponds to the fraction of fusion peaks that are contained in the overlap with each ChIP-Atlas cell type. Color is based on p value determined empirically by computing the overlap between random BED files and each ChIP-Atlas cell type. Only fusions with >40 differential peaks compared to EV and significant overlap (-log p value > 3 and fraction > 0.2) with at least one ChIP-Atlas cell type are shown and the kinase fusions are omitted from the visualization.

To examine mechanisms across all variants in the library, we looked for DNA motifs that were enriched in each variant’s differentially accessible peaks. In all cases where a fusion contained a known DNA binding domain (DBD), the motif associated with that DBD was highly enriched in peaks with increased accessibility for that fusion (Fig 4c,d; Supp Fig 5). For instance, peaks induced by all fusions containing NR4A3 had enrichment of the Nur77 motif (a nuclear receptor 4A family protein), peaks from all fusions with ATF1 had enrichment of the ATF1 motif, and the most significantly enriched motif in EWSR1-NFATC2 regulated peaks was NFAT. The most significantly enriched motif for nuclei expressing EWSR1-FLI1 was the FLI1 binding motif GGAA (Fig 4c,d) and indeed EWSR1-FLI1 marker peaks have significantly higher content of GGAA repeats compared to marker peaks from control cell lines (Supp Fig 3d) consistent with previous data^44–46^. In one case, the enriched DNA motif was unexpected; we found that IRF2BP2-CDX1 pioneered chromatin at loci enriched for the p53 motif (Fig 4d). IRF2BP2-CDX1 has only been reported once and never mechanistically studied. However, IRF2BP2 overexpression alone has been shown to antagonize p53-induced activation of cell cycle and pro-apoptosis genes. That IRF2BP2-CDX1 increases accessibility at p53 loci suggests either that IRF2BP2’s fusion to CDX1 alters its interaction with p53 or that this interaction is not well modeled in 293T cells. In general, we find substantial concordance between fusion pioneering and their respective domain components even for rare fusions.

Interestingly, we find striking chromatin accessibility changes even for fusions that are seemingly unable to directly bind DNA. Despite not having DBDs or known direct interaction with DNA, differential peaks for all 4 kinase fusions were heavily enriched in the Fos/Jun motif. ETV6-NTRK3 has been previously shown to activate the AP-1 complex^47^. This suggests that CCDC6-RET, FGFR3-TACC3, and TPR-NTRK1 likely activate the AP-1 complex in a similar fashion to ETV6-NTRK3. Similarly, EPC1-PHF1 and MEAF6-PHF1 lack DNA binding domains but have significant enrichment of the Tcf4 motif at differentially accessible peaks. Tcf4 is a key mediator of Wnt signaling and both fusions cause low grade endometrial stromal sarcoma (LGESS) which has been shown to have increased Wnt signaling^48^. Our data suggests that Wnt activation in LGESS is directly due to fusion expression and is true for both of these causal fusions. Taken together, PROD-ATAC is therefore capable of resolving both direct and indirect modes of chromatin dysregulation.

Finally, we sought to determine whether signatures of cancer and cell-type specificity could be recovered from the fusion-produced chromatin alterations in 293T cells. To this end, we examined the overlaps between peaks regulated by fusions in 293T and signature ATAC peaks across thousands of known human cell types. For each fusion, we counted the number of overlaps between that fusion’s differentially accessible peaks (compared to EV control; Supp data 1) and those ATAC peaks of known human cell types from ChIP-Atlas. To determine the significance of the overlap, we used an empirical null distribution created by permuting the chromosomes within the list of fusion-specific peaks and recalculating the overlap with the same known cell types (Supp data 2). Peaks defined for the four kinase-containing fusions overlapped with hundreds of cell types without an obvious pattern possibly owing to their aberrant activation of AP-1 complex (Fig 4d). Subsetting to the non-kinase pioneer fusions revealed that many fusion-regulated peaks are shared with cell types relevant either to the cancer the fusion causes or to biological processes that the fusions’ components are known to regulate (Fig 4e). For instance, PAX7-FOXO1 peaks strongly overlapped with peaks from alveolar rhabdomyosarcoma (RMS) cell lines including RMS13, Rh4, and Rh30, which was also seen quantitatively at several loci in pseudobulk (Supp Fig 4b). Interestingly, the peaks from the rare fusion ACTB-GLI1 overlapped with some of the same RMS cell types. ACTB-GLI1 has not been seen in RMS, but it is exceptionally rare making a lack of association with this cancer type underpowered. Interestingly, *GLI1* is recurrently amplified in RMS^49^ and *GLI1* upregulation has been found in drug-resistant RMS^50^ suggesting a possible shared mechanism between PAX7-FOXO1, RMS, and this rare fusion. Together, this shows the fascinating feature of many driver fusions that their effects are context invariant and that seemingly cell-type specific knowledge can be recovered even from 293T cells.

Several additional known cell-type specific relationships appeared suggesting that many oncogenic mechanisms can be inferred even from 293T cells. This included a significant overlap between EWSR1-FLI1 peaks and the Ewing sarcoma cell line A-673, overlap between TMPRSS2-ERG and human umbilical vein endothelial cells (ERG is a canonical vascular endothelial cell regulator^51^), and overlap between TMPRSS2-ERG and EWSR1-NFATC2 peaks with those of osteoclast precursor cells (both fusions regulate cancers that cause osteolytic lesions^52^ or otherwise disrupt the osteoclast:osteoblast balance^53^). Although not displayed, peaks from both EPC1-PHF1 and MEAF6-PHF1 which cause endometrial stromal sarcoma had highly significant overlaps with ATAC peaks from endometrial stromal cells (p = 0.001 and p=0.005, respectively, two-tailed t-test; Supp data 2). Interestingly, peaks regulated by ARFGEF2-HNF4A strongly overlap with those from gastrointestinal cell types including the liver and colon. HNF4A is a transcription factor that plays an important role in the specification of GI cell types^54^ and HNF4A amplification has been found in colorectal cancer^55^. While there are no publications of ARFGEF2-HNF4A and it has only been reported once in TCGA in an unspecified sarcoma^56^, this data would suggest that ARFGEF2-HNF4A might be altering chromatin at loci consistent with HNF4A’s known role in lineage specification during GI cell type development. These data collectively show the surprising result that many oncofusions have context-invariant effects which are recovered by PROD-ATAC, and that it may be possible to learn certain cell-type relevant mechanisms even in a cancer-agnostic background.

### Some oncofusions have gain-of-function pioneer abilities not attributable to their constituent parts

While some oncofusions exhibit gain-of-function effects compared to their individual domains^18,57^, it is not clear how widespread this property and what features govern whether or not a given fusion exhibits gain-of-functionality. Similarly, it is unclear whether fusions that share one (but not both) domains generally exert the same or different effects on chromatin. PROD-ATAC offers a large-scale approach to investigate these questions by examining the function of individual domains, full length wild-type proteins from which domains are derived, and their combined fusion products concurrently. To test these hypotheses, several controls for many of the most highly recurrent fusions and commonly involved domains were included in the library. This included several full length wild-type proteins from which part of recurrent fusions are derived including ATF1, NR4A3, DDIT3, EWSR1, and FLI1. We also included several domains alone to test if their overexpression alters cell-state even without the remaining component (whether from the fusion or the wild-type protein). These include EWSR1, FUS, ATF1, NTRK3, and ETV6. In some cases, gain-of-function features were strong enough that they were visible in the distribution of cells in UMAP space. For instance, nuclei containing the full length wild-type ATF1 protein largely overlap the distribution of cells that contain the empty vector control whereas cells expressing either EWSR1-ATF1 or EWSR1-CREB1 occupy a *de novo* cell state (Fig 5a, left). Similarly, cells expressing the full length NR4A3 wild-type protein overlap with cells expressing the empty vector control but are clearly distinguishable from those expressing the TAF15-NR4A3 fusion (Fig 5a, right). These gain of function features are also clear when examining marker peaks for each of these fusions (Fig 5b). EWSR1-ATF1, EWSR1-CREB1, and FUS-ATF1 have many similarities across thousands of peaks and almost none of the marker peaks for these three fusions are recapitulated with either the relevant full length wild-type or domain controls.

**Figure 5:**
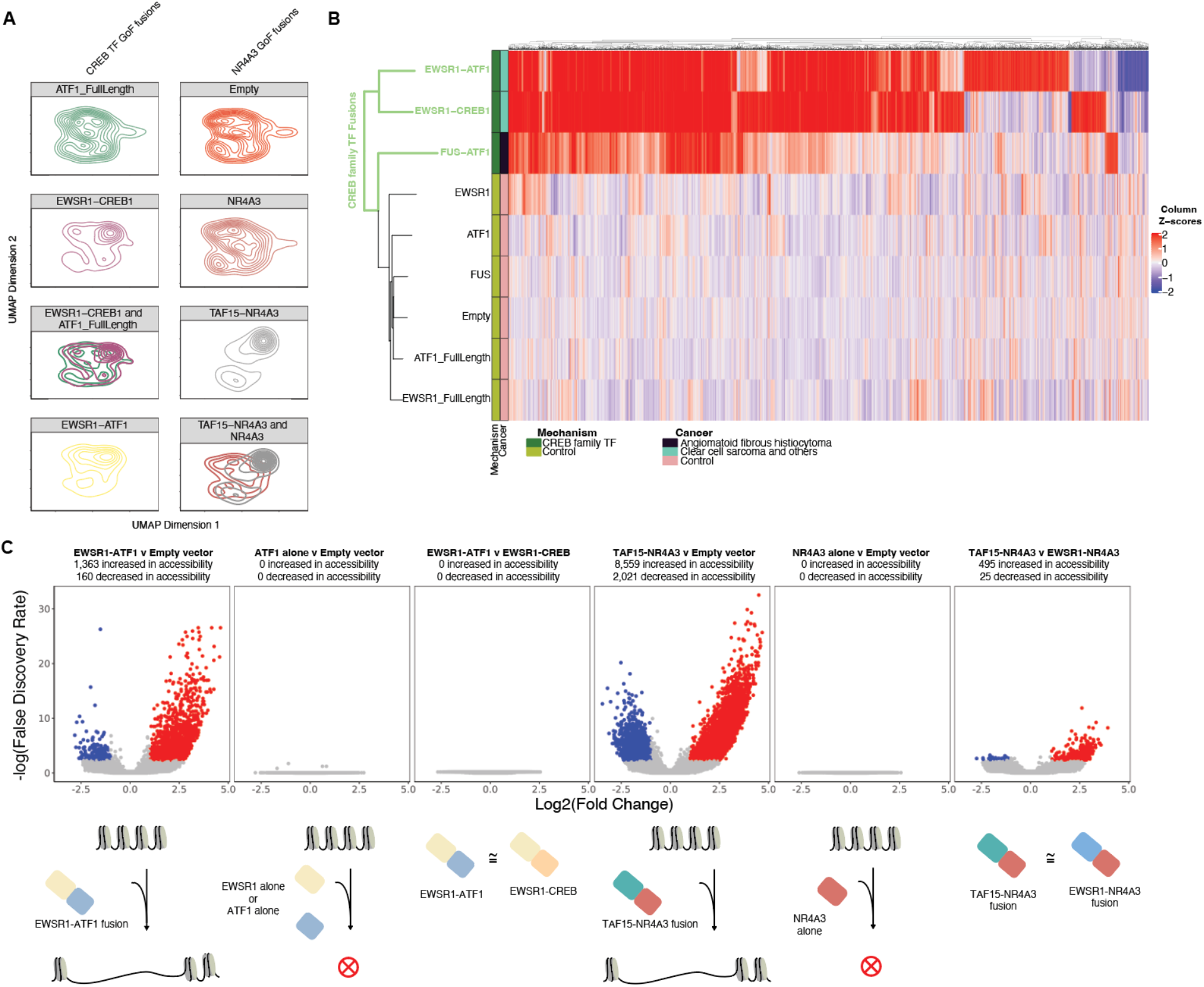
Some oncofusions have gain of function pioneer activity not attributable to individual components. (A) Density contours of UMAP embedding after latent semantic indexing with individual variants highlighted. Cells expressing EWSR1-ATF1 and EWSR1-CREB1 have an altered distribution relative to the full length ATF1 control as do cells expressing TAF15-NR4A3 relative to those expressing the full length NR4A3 control. (B) Heatmap of z-score normalized accessibility at 4,856 peaks with |z-score| > 1.5 for at least one of EWSR1-ATF1, EWSR1-CREB1, or FUS-ATF1. (C) Representative volcano plots of -log(false discover rate) versus log_2_(fold change) for peaks comparing fusions to relevant domain controls. Red peaks are increased in accessibility whereas blue peaks are decreased in accessibility.

We next systematically tested for gain-of-function pioneer activity among many previously never studied fusions. To this end, we made all relevant comparisons of pseudobulk replicates among individual domains and fusions. For example, while EWSR1-ATF1 significantly increased accessibility of over 1,000 peaks compared to empty vector control, there were 0 peaks that were differentially accessible when the full length wild-type ATF1 protein was expressed (Fig 6c). This is particularly interesting because EWSR1-ATF1 loci are highly enriched in the ATF1 motif (Fig 4c), suggesting that the EWSR1 N-terminus (but not the wild-type ATF1 N-terminus) is sufficient for inducing pioneering at ATF1 sites. Similarly, while TAF15-NR4A3 increased accessibility at almost 8,600 peaks within which the NR4A family motif was highly enriched (Fig 4c), expression of the full length NR4A3 wild-type protein alone changed 0 peaks. We also examined whether gain-of-functionality and chromatin pioneering for a given fusion depends on the fusion partner. When directly compared, EWSR1-ATF1 and EWSR1-CREB1 had no significantly different peaks whereas the direct comparison of TAF15-NR4A3 and EWSR1-NR4A3 revealed only a few hundred altered peaks out of the 8,600 total pioneered. Generally, there were significantly fewer significant peaks when comparing fusions of the same class than when comparing fusions to controls (Supp Fig 6). This suggests that DNA binding is unsurprisingly directed by the DBD but, interestingly, that pioneering is further orchestrated by the partnering domain in ways often not recapitulated by the wild type proteins. These are striking gain-of-function effects and while such GoF has been shown for EWSR1-FLI1, it has not been shown for EWSR1-ATF1, EWSR1-CREB1, TAF15-NR4A3, TFG-NR4A3, EWSR1-NR4A3, or TCF12-NR4A3 all of which are shown here. Our data suggests that gain-of-function chromatin pioneering is a common feature shared among many oncofusion proteins.

**Figure 6:**
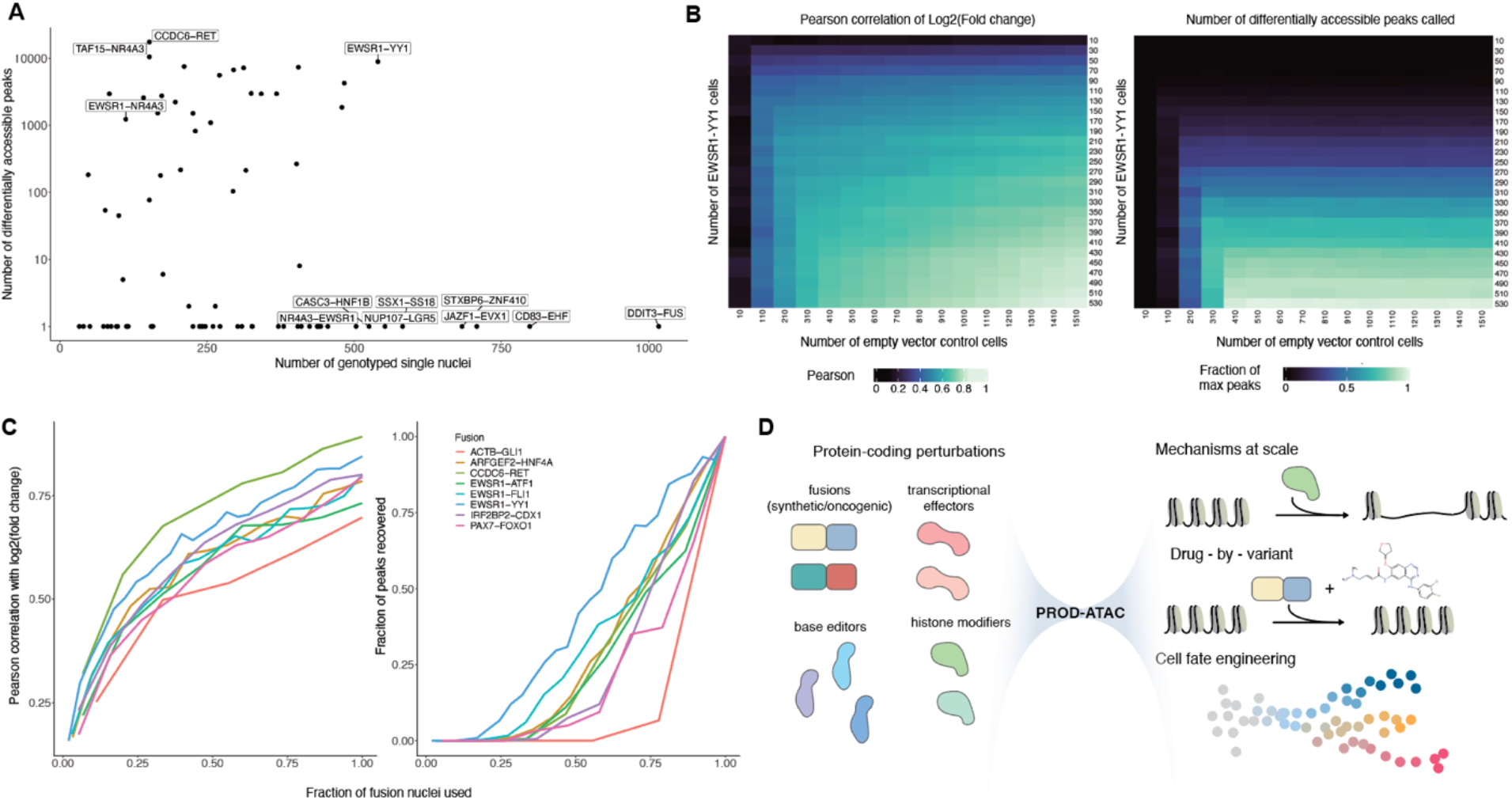
Down-sampling demonstrates that PROD-ATAC is a scalable method to discover mechanisms for hundreds of protein-coding variants simultaneously. (A) Number of significantly differentially accessible loci for each variant compared to empty vector control versus the number of genotyped cells with Spear-ATAC. (B) Down-sampling the number of empty vector control cells and EWSR1-YY1 expressing cells. Pearson correlation of log_2_(fold change) between fusion and control cells for all 37,701 peaks (left) and number of peaks called as differentially accessible after down-sampling (right). (C) Down-sampling fusion-containing cells for 8 variants while maintaining constant number of empty vector control cells evaluated by Pearson correlation of log_2_(fold change) values (left) and number of recovered differentially accessible peaks (right). (D) Future applications of PROD-ATAC to learn disease mechanisms, identify drugs that target specific variants and mechanisms, and evaluate the mechanisms of synthetic factors for directing user-defined cell states.

### Discovering genetic mechanisms and identifying modifiers at scale

The power of PROD-ATAC is scalability and generalizability. PROD-ATAC can be scaled to measure the effects of many hundreds to thousands of genetic variants simultaneously. Besides oncofusions, libraries could contain missense variants in transcription factors, chromatin readers and writers, signaling effectors, and even structural proteins that alter chromatin architecture; short ORFs or microproteins of unknown function; ancestral protein variants throughout the tree of life; or synthetically designed proteins. A key determinant of the scalability of PROD-ATAC is the number of nuclei needed to resolve each variant’s effects. This depends on the number of loci regulated by the variant, the variant’s effect size at each locus, and the likelihood of capturing that information given the known biases of Tn5. To guide future larger scale experiments, we evaluated the parameter-space required to resolve variant effects by progressively down-sampling the number of nuclei captured for many variants.

We captured many hundreds of nuclei per genotype (average 331, interquartile range 172 – 437 after filtering) and we resolved variant effects with as few as ∼100 nuclei per variant (Fig 6a). We also deeply sequenced the chromatin libraries resulting in more than 56,000 high-quality fragments per nucleus on average (interquartile range 48,769 – 61,911 after filtering). The information content for each genotype on average was therefore comparable to that of a typical bulk ATAC-seq experiment with tens of millions of reads per condition, which allowed us to resolve a wide range of effects. For instance, we detected that CCDC6-RET regulated nearly 20,000 differentially accessible peaks after having captured only 152 nuclei. Similarly, we detected 1,235 differentially accessible peaks when expressing EWSR1-NR4A3 across only 112 nuclei, yet we detected 0 peaks for the reciprocal control NR4A3-EWSR1 when averaging over 503 nuclei. That the false discovery rate is well-calibrated suggests that false positives are unlikely to be seriously problematic as the scale increases.

To determine how sensitive our measurements are to changes in the number of nuclei captured, we down-sampled the number of nuclei for several variants that exert a wide range of effects. In the case of EWSR1-YY1 (one of the strongest pioneer fusions in our library), the Pearson correlation between the down-sampled and full-scale data is robust to reducing either the number of EWSR1-YY1 nuclei or the number of empty vector control nuclei (Fig 6b, left). While we captured 1,510 empty vector control nuclei, only a few hundred are necessary to retain a high correlation coefficient. Similarly, the number of EWSR1-YY1 containing nuclei can be down-sampled to as few as ∼300 nuclei (∼600 nuclei originally) while retaining a high Pearson correlation coefficient (∼0.8). In general, only a few hundred nuclei per genotype are required to resolve variant effects, which is comparable in information content to standard bulk ATAC-seq experiments.

While the Perason correlation of fold changes for all peaks is robust to down-sampling, the number of recovered differentially accessible peaks (a metric of higher resolution) is significantly more sensitive. This is similar for fusion and empty vector containing nuclei both of which require many hundreds of nuclei to retrieve a large fraction of known differentially accessible peaks (Fig 6b, right). We tested the generality of this property by fixing the number of control empty vector cells and down-sampling fusion containing cells for 8 different fusions. These fusions had a wide range of effect sizes and mechanisms. In each case, the pattern held that the correlation with the original data is more robust to down-sampling than is the number of differentially accessible peaks retrieved (Fig 6c). Those variants with the most negative slopes were those with the fewest nuclei captured in total suggesting that we likely have not achieved saturation for these variant’s effects. Together, these analyses suggest that significant information could be retained with only a few hundred nuclei per library member meaning that PROD-ATAC could be applied to test hundreds to thousands of variant effects simultaneously. Variants with complex features and for which saturation is not achieved in the first experiment could be identified quickly with low nuclei counts and then subject to deeper phenotyping in future experiments.

## Discussion

PROD-ATAC is a scalable method for interrogating genome-wide chromatin dysregulation induced by arbitrary sets of protein-coding variants. PROD-ATAC allows researchers to systematically explore the impacts of hundreds to thousands of protein-coding variants—without requiring any prior knowledge of variant effect—to learn both mechanisms of and relationships between variants in a relatively unbiased way. To our knowledge, no prior high-throughput experiments have examined the sufficiency of causal variants to disrupt chromatin for hundreds of variants simultaneously and outside of their native contexts. That expression of causal variants in non-native contexts can report on cell-type relevant mechanisms is fascinating and should promote future high-throughput screens in unified model systems like this.

We used PROD-ATAC to produce a global view of the landscape of fusion-induced chromatin disruption. In most cases, the structure revealed was consistent with known prior mechanisms. For instance, clusters based on chromatin accessibility clearly discriminated among ETS family fusions, kinase-containing fusions, NR4A3-containing fusions, and CREB-family transcription factor containing fusions. However, we also learned non-intuitive relationships that would’ve otherwise been difficult to predict *a priori* or to learn with existing low-throughput reverse genetics methods. In some cases, we see evidence of convergent mechanisms among seemingly disparate fusions. For instance, the rare fusion ACTB-GLI1 (which causes mesenchymal tumors) and the relatively common fusion PAX7-FOXO1 (which causes rhabdomyosarcoma) converge on similar chromatin disruption (and indeed *GLI1* amplification and overexpression is seen in rhabdomyosarcoma). In other cases, we see heterogeneity even among fusions that cause the same cancer types. For example, several fusions cause synovial sarcoma. While we saw no evidence of pioneer activity among the most common variants (SS18-SSX1/2/4) in 293T cells, the very rare synovial sarcoma fusion, SS18L1-SSX1, did cause strong pioneering. This global view of variant mechanisms—spanning rare and common mechanisms across dozens of cancer types—is possible with PROD-ATAC.

We also showed the interesting result that many causal factors exert context-invariant effects. Variant effects recovered in 293T cells can even allude to each variant’s context of origin. For instance, EWSR1 fused to ETS-family proteins all pioneered chromatin at loci shared with Ewing sarcoma cell lines. PAX7-FOXO1 pioneered chromatin at loci that overlap with several rhabdomyosarcoma cell lines as did EWSR1-ATF1 at clear cell sarcoma derived peaks. We recapitulated several known biochemical features of oncofusions as well including that FUS-DDIT3 and EWSR1-FLI1 exhibit gain-of-functionality. No doubt *in vitro* model systems cannot recapitulate all context-specific variant effects, and in some cases we identified possible inconsistencies between the 293T and cancer-specific models (e.g. IRF2BP2-CDX1 increasing accessibility at loci with the p53 motif). While inaccuracies in cancer models can be problematic, cases of discordant findings lead to important hypotheses about context-specific mechanisms. In this way, having a unified context to study variant effects is not only a worthwhile tradeoff to increase throughput and to unambiguously identify causal effects, but it also allows researchers to probe context-specific effects. This is impossible in the complex milieu of patient samples and with low-throughput methods. However, this type of discordance was rare, and we showed striking recapitulation of known or suspected mechanisms of action across dozens of unique fusions representing dozens of cancer types. Taken together, this suggests that oncofusions may be sufficient (independent of context) for inducing oncogenic cell-state changes, a reasonable hypothesis given that fusion-positive sarcomas often have low mutational burdens^15,41,42^.

Rare and orphan variants are unlikely to be investigated if not for high-throughput methods like PROD-ATAC, and such variants represent patients who could benefit from having their oncogenic drivers studied. Population-scale sequencing efforts have revealed a preponderance of rare germline variants nearly half of which are singletons^58^; a similar growing list of rare variants exists in cancer biology as more tumors have been sequenced. It is critical to identify the effects even of the rarest of variants. That biochemical features of oncofusions are robustly modeled in the 293T context allowed us to hypothesize about mechanisms even of rare fusions some of which have never been studied. Studying common *and* rare genetic variation will allow researchers to identify convergent mechanisms of disease that would be otherwise hidden from studies that focus only on common variants. Convergent mechanisms across the variant frequency spectrum are also more likely to be of biological significance. Systematically dissecting the landscape of variant effects with methods like PROD-ATAC will reveal more complete and compelling pictures of variant effects and allow researchers to identify a high-quality set of mechanisms that underly complex biology for therapeutic targeting.

PROD-ATAC is a powerful hypothesis-generating tool for mechanistic biochemistry studies. Based on our findings, we hypothesize that phase separation is a recurrent feature of oncofusions and might underly their ability to pioneer chromatin. For example, we showed that many NR4A3 containing variants have gain of function pioneering regardless of the N-terminal domain. Most of the N-terminal domains in these cases contain long stretches of prion-like domains (PLDs)^57,59,60^, and PLDs are frequently implicated in phase separation^61,62^. This result is important given that a high-throughput screen exists for dissolving phase separated oncogenic condensates^63^. That the N-terminal component drives a biophysical property (i.e. phase separation) could also explain why these fusions induce similar chromatin alterations despite having seemingly diverse N-terminal domains. PROD-ATAC allows researchers to hypothesize about these types of effects at scale and learn how individual domain components modulate fusion activity. Experiments to definitively answer these questions would require several dozens of controls per fusion (individual domains, full length wild type proteins, all N-C terminal pairwise combinations swapped, etc.) which are intractable with current low-throughput methods.

We anticipate several exciting applications of PROD-ATAC for the discovery of genetic mechanisms at scale (Fig 6d). First, questions regarding the context-specific nature of protein-coding variants can only be determined unequivocally in well-controlled settings like these. Sequencing primary samples is critical, but causality will be difficult to ascertain without controlled perturbation-based experiments. Using this method across many contexts (e.g. creating landing pad acceptor cells in various contexts) will allow researchers to ask how concomitant changes to cellular and genetic background impact hundreds of coding variants of interest simultaneously. Second, because biochemical mechanisms are well-modeled, we anticipate that PROD-ATAC will be useful for screening libraries of small molecules or libraries of additional genetic perturbations that rescue or synergize with variant effects (e.g. CRISPR libraries *combined* with protein-coding libraries). That is, rather than be relegated to low-resolution screens based on cell growth or single-gene reporter fluorescence, researchers can use PROD-ATAC to identify modifiers of high-dimensional cell states. Of course, chromatin accessibility is not the only important complex cell state. We anticipate perturbation experiments with multi-omic measurements will be useful for creating networks of regulation altered by thousands of perturbations simultaneously. Combining several modalities of high-content readouts with libraries of coding variants will complete the variant-to-function pipeline from chromatin regulation to gene expression to cellular morphology and behavior. Finally, PROD-ATAC imposes no limitations on the types of protein-coding variants examined. This will be a useful tool for assaying libraries of both naturally occurring variants and synthetic variants towards engineering cell states and fates.

## Methods

### Cell culture

293T cells were a gift from Peter Lewis’s laboratory who previously purchased them from ATCC. Cells were validated by STR analysis and regularly confirmed to be mycoplasma free by Venor GeM Mycoplasma Detection Kit (Milipore Sigma). 293T cells were cultured at 37 C; 5% CO_2_; in high glucose DMEM supplemented with 10% tet-system approved FBS (Gibco), 1% penicillin-streptomycin, and 1% GlutaMAX (Gibco); and they were routinely passaged before achieving confluence.

### Lentivirus production and landing pad generation

We created a clonal line of 293T cells that harbored a single-copy of a previously published landing pad for site-specific, irreversible library recombination. The attP-containing landing pad and several attB-containing control donor plasmids were gifts from Kenneth Matreyek. The landing pad used (pLenti-TetBxb1-BFP-rEF1a-rtTA3G, LLP-Growth) allows for inducible expression of recombined libraries and selection of recombined cells that is separate from selection of cells containing the landing pad itself. Lentivirus containing the landing pad sequence was produced by co-transfecting 293T cells at ∼50% confluence with psVSV-G (Addgene #12259), psPAX2 (Addgene #12260), and LLP-Growth plasmid using lipofectamine 3000 at a ratio of 1 : 3.5 : 3.5, respectively. Supernatant containing virus was collected 48 hours after transfection, centrifuged at 500 × g for 10 min, filtered through an 0.45 *μ*m PES filter, and used immediately. To create the landing pad containing cell line, 293T cells in 6 well dishes at ∼50% confluence were transduced with either 0, 100 *μ*L, 500 *μ*L, 1 mL, or 2 mL of viral supernatant. Replicate wells were induced with 2,000 ng/mL doxycycline at the same time of transduction and a separate well (that was seeded at the same time as all others) was trypsinized and counted to later estimate multiplicity of infection (MoI). 48 hours after induction the percent blue fluorescence protein (BFP) positive (BFP^+^) cells was determined by flow cytometry. Conditions with an estimated MoI << 0.1 were retained and successfully transduced cells were selected for with the addition of 6 *μ*g/mL blasticidin for 10 days.

We then generated a clonal line of landing-pad containing cells from the condition that was initially transduced with 100 *μ*L virus; we chose this condition because it was the smallest amount of virus for which there were appreciable transductants (and therefore very few of these cells would contain multiple integrated landing pad sequences). After selecting with blasticidin, we induced expression of BFP with the addition of doxycycline and sorted single cells into wells of a 96 well plate. Blasticidin was added to all media for these cells in perpetuity. Individual cells were allowed to grow for approximately 1 week in media containing 20% FBS before being expanded and cultured under normal conditions. To determine those clones with the widest on and off state for transgene induction, we induced all outgrown clones with and without doxycycline and used flow cytometry to measure BFP expression. The clone 293T-B4 was chosen given that it had the lowest variance in BFP expression with doxycycline and least amount of leaky BFP expression without doxycycline (Supp Fig 1a). This clone was validated to contain a single copy of the landing pad transgene by two methods. First, we recombined a library containing equal parts attB-mCherry and attB-GFP. The lack of double positive cells suggested only a single copy of the landing pad was present (Supp Fig 1b). Second, we performed droplet digital PCR with primers for the landing pad sequencing and the housekeeping gene *ALB*. That the copy number of the landing pad was approximately half that of the endogenous *ALB* control also suggested that only a single copy is present (Supp Fig 1c). The 293T-B4 clonal line was used for creating all libraries in this work.

### *In silico* oncofusion library curation

A list of fusions was downloaded from COSMIC^34^ in October 2022. 3 distinct sets of fusions were considered; (1) any fusion in the COSMIC database with “sarcoma” present in the primary histology field (n=55), (2) known sarcoma fusions (n=76) derived from Table 1 of Perry et al. *Annu. Rev. Cancer Biol.* **2019** and from ChimerDB v4.0^20^ (with the requirement that they were in-frame fusions, present ≥ 5 times, contained a known transcription factor domain, and had a Seq+ annotation), and (3) a list of 12 fusions not derived from sarcomas (e.g. TMPRSS2-ERG, TPR-NTRK1, CCDC6-RET, FGFR3-TACC3, PML-RARA, BCR-ABL1, BCR-JAK2, etc.). This resulted in 105 unique fusions considered further as 38 fusions were both in the known sarcoma fusions and COSMIC database with sarcoma in histology. We further restricted this list by only considering those fusions found in the COSMIC database. For each unique pair of 5’ and 3’ genes fused, the COSMIC ID for the most common variant (as assessed by number of unique sample IDs in fusion database) was considered. For this fusion ID, for the 5’ (respectively, 3’) gene, the chromosome, strand, last observed exon (first observed exon), genome start from, and genome stop from were extracted. Nucleotide sequences for relevant transcripts were found in GENCODE v28^64^ and the coding sequence for the relevant parts of the gene included in the fusion were extracted, resulting in 2 nucleotide sequences corresponding to the sequence encoding the N terminal and C terminal domains separately. If the total length of the combined nucleotide sequences was divisible by 3, this sequence was finalized. If the combined sequence was not divisible by 3, each individual sequence was shortened by 1 or 2 base pairs to make it divisible by 3 to avoid introducing a frameshift. The amino acid sequences derived from human tumor samples for several well-known fusions are present in GenBank. In these cases, we confirmed that the sequences we mined matched those previously published. For instance, our SS18-SSX1 sequence was a 100% match with BAF56182.1, our PAX3-FOXO1 sequence was a 100% match with AAC50053.1, our SS18-SSX4 sequence was a 99.6% match with AAG31034.1, and our CCDC6-RET sequence was a 100% match with BAM36435.1. Our EWSR1-FL1 sequence matched entirely to ACA62796.1 and was a 98.8% match with ADX41459.1. The difference between these two references is that the latter is missing 6 amino acids in the EWSR1 domain. The most common EWSR1 isoform lacks these 6 amino acids whereas some other isoforms include them. Fusions in patients have been found using both isoforms and because we used the most common isoform for all fusions, our EWSR1 sequences across all fusions lack these 6 amino acids. This is the same reason that our EWSR1-ATF1 sequence is a 98.9% match with ADX41457.1, again differing only by those 6 amino acids. Overall, this shows that our method of generating fusion sequences was highly concordant with what is found in patient samples for those that are published.

In addition to the fusions, several controls were added to the library. Several of the most common domains from the fusion list (e.g. EWSR1, ATF1, FUS, FLI1, ETV6, NTRK3 etc.) were added without their fusion partner. Next, for several of the most common components we additionally included the full length wild-type counterpart (e.g. EWSR1, FLI1, ATF1, DDIT3, NR4A3 etc.). Sequences were then filtered to be less than 4.8 kb in total. Finally, an empty vector control sequence was added that did not contain an open reading frame. This yielded 116 total unique sequences. To help promote even expression across library members, ATG was added to initiate the open reading frame for all those sequences that did not already have a start (i.e. domain controls derived from the C terminal half of a fusion protein) with the exception of the empty vector control. Barcodes were added to these sequences (see Methods: Plasmid library construction and sequencing). The final sequences were ordered from Twist Biosciences as clonal genes and only two were unable to be synthesized (EWSR1-ERG and NR4A3-TAF15).

### Variant barcoding scheme

In Spear-ATAC, each nuclei needs to be associated with its expressed guide RNA (gRNA) perturbation^14^. Because gRNAs are small, Spear-ATAC directly reads out the gRNA from the enriched library for each nucleus. To make our method amenable to arbitrary protein-coding perturbations, some of which can be several kilobases long, we modified this scheme to include short DNA barcodes to represent each protein-coding perturbation. In this case, each protein-coding member of the library contains two barcodes (Supp Fig 1d). First, there is a hardcoded 12 nucleotide (nt) DNA barcode that is synthesized with each variant coding sequence. The DNABarcodes package in R was used to generate a list of 200 barcodes each that is 12 nt with minimum hamming distance of 5 and is GC balanced. Each member of the variant library was then randomly assigned to one of these molecular barcodes and that assignment was retained in perpetuity. Ideally, this barcode would be flanked by Nextera adapters such that it could be identified after enrichment from the single-cell library (like the gRNA is flanked in Spear-ATAC). In this hypothetical situation, only one barcode is needed. However, many commercial oligonucleotide synthesis companies disallow the inclusion of Nextera or TruSeq adapters internally given that they are also used for quality control with next-generation sequencing (NGS). To circumvent this, we added a second barcode of 16 random nucleotides between Nextera adapters that is cloned into a position adjacent to the hardcoded barcode. The random barcodes were synthesized as N_16_ within primers ordered from Integrated DNA Technologies. The result is that the random barcode and hardcoded barcode are each within113 nt of each other. This allowed us to sequence the known (hardcoded) and random barcode together in one short-read sequencing run to create a transitive mapping between each variant, hardcoded barcode (known *a priori*), and the random barcode. When a particular random barcode was seen after Spear-ATAC, we would know (based on the short-read sequencing map) which hardcoded barcode it is associated with and therefore which variant is present for that nucleus. Having both barcodes therefore generalizes this format to any possible method of synthesizing the library of variants. This scheme also allowed us to quantify template switching and barcode hopping between hardcoded and random barcodes after recombination, which occurred exceptionally rarely (23 of 8,547, or 0.27%, random barcodes switched which hardcoded barcode they were assigned to after Bxb1-mediated recombination). The complete list of variant nucleotide sequences, expected protein product, and hardcoded barcodes is given in Table 1.

### Plasmid library construction and sequencing

All oncofusion and control sequences were prepared for downstream one-pot BsmBI-based Golden Gate Assembly into an attB-containing donor plasmid for ultimate recombination into attP-containing 293T landing pad cell lines. This included codon-optimizing all open reading frames for human expression (with Twist Bioscience online software), removing internal BsmBI sites by creating synonymous substitutions that avoid rare codons, appending a C terminal G-A-G linker and HA epitope tag to all coding sequencing, adding the proper hardcoded barcode sequence as described above, and flanking this cassette by universal primer binding sites such that all library members could be amplified with a single primer pair. This primer pair notably includes part of the Nextera adapter sequences and the oSP2053 primer binding site needed for Spear-ATAC. The primers also contain BsmBI sites needed for Golden Gate Assembly with the attB-containing backbone (see below). All library members were ordered from Twist Bioscience and then amplified by PCR. Of the 114 unique sequences, 113 were amplified with one clearly dominant product at the correct size by agarose electrophoresis. PCR products for these 113 variants were quantified using the AccuClear Ultra High Sensitivity dsDNA Quantitation Kit and then pooled based on their sizes to ideally achieve a uniform distribution.

At the same time, the attB-containing backbone was linearized by PCR and prepared to accept the library inserts via BsmBI Golden Gate Assembly. This PCR contained primers that linearized the backbone and added an N_16_ random barcode, part of the flanking Nextera adapters (the other part was supplied by primers used in the PCR that amplified the insert library), and BsmBI sites. The backbone PCR was followed by DpnI digest by spiking in 2 uL DpnI, 10 uL NEB rCutSmart buffer, and water to reach a total volume of 100 uL. The reaction proceeded at 37C for 1 hour followed by a 20 min inactivation at 80C before being cleaned up. A single Golden Gate Assembly reaction was then performed to insert the pool of variant members into the universal attB-containing backbone. This was performed by combining 150 ng of linearized, DpnI digested, and cleaned up backbone with cleaned up variant library PCR product for an approximately 2:1 molar ratio (the insert sizes varied because library member sizes vary, so this calculation was based on the median size). To this was added 2 uL T4 buffer, 1 uL BsmBI-v2, and water to bring the total reaction to 20 uL. The reaction proceeded at 42 C for 1 hour followed by 60 C for 5 minutes. The 20 uL product was dialyzed on a MF-Millipore Membrane Filter (0.025 um pore) for 1 hour with a dialysate of double distilled water.

From there, a bacterial stock of the library was generated. 2 uL of the dialyzed golden-gate product was transformed into 25 uL of electrocompetent DH10B E coli cells using a Bio-rad MicroPulser (165-2100), Ec2 setting (2 mm cuvette, 2.5 kV, one pulse). Two replicates of this procedure were performed. Cells were immediately recovered in 973 uL of pre-warmed SOC while shaking at 37 C for 1 hour. Critically, the transformation efficiency determines not only library coverage but also the number of expected random barcodes (which are thought to be in significant excess after Golden Gate compared to the number of successful transformants). To ensure that each library member receives several random barcodes (for redundancy) and to ensure that the number of random barcodes does not swamp the number of expected nuclei captured with single-cell sequencing (i.e. so that we would sequence each random barcode several times across several nuclei), we throttled the number of transformants retained. To determine the number of transformants to retain, we first created spot plates from each of the replicates. Each spot contained 2 uL of culture and we spotted undiluted, 1:10 diluted, and 1:100 diluted spots each in triplicate. These plates grew overnight at 37 C. At the same time, we split the two 1 mL liquid cultures into several subsets: 100 uL and 700 uL from replicate 1 and 50 uL, 100 uL, and 700 uL from replicate 2. To each of these, LB and carbenicillin was added and the cultures were grown overnight while shaking at 37 C. After overnight growth, the transformation efficiency was estimated based on the spot plate in both replicates to be ∼150-220 colonies per variant. This meant that the 100 uL sample of the initial recovered culture would contain an appropriate number of barcodes. Replicate 1 with 100 uL was therefore plasmid prepped and a glycerol stock (25% v/v sterile glycerol stored at −80 C) was simultaneously prepared.

We sequenced the random barcode and the hardcoded barcode after insertion into the plasmid attB-containing backbone with 1 × 150 bp reads. To filter for reads that contain the correct structure of amplicon, we grepped for 4 constants sequences that flanked the barcodes: “AAATCCAAGC”, “CCAGAGCATG”, “CAAGGTGGTT”, and “ATACTGATTC”. This yielded 5,625,954 amplicons to analyze. We then matched each of the supposed hardcoded barcodes to one of the known hardcoded barcodes from the list that we intended to synthesize. This allowed for synthesis, sequencing, or PCR errors that would slightly alter the read-out hardcoded barcode compared to what the Twist order intended. For all but 4, we were able to identify a closest match using a maximum restricted Damerau-Levenshtein distance of 3. Each hardcoded barcode was assigned to its closest match and the 4 unmatched barcodes were discarded. We did not have such a white-list of the random barcodes and therefore could not do the same matching procedure for these. Instead, we first generated a list of unique hardcoded barcode – random barcode pairs of which there were 48,582 sequences that were approximately evenly distributed about the expected frequency except for a fat tail of rare pairs. Restricting that list to pairs to those that were seen more than once yielded 14,803 unique pairs. Finally, we required that, for a given random barcode, more than 95% of the reads had to match to the same hardcoded barcode. That is, should a random barcode frequently map to more than one hardcoded barcode, we would not be able to confidently assign it to a particular perturbation. This criteria further restricted the pool yielding 12,269 confident barcode pairings with which we can ultimately assign each nuclei to a perturbation identity. There was a strong correlation between the distribution of confidently assigned random barcodes inserted into genomic DNA and those in the initial plasmid library (Supp Fig 1g).

### Library recombination and sequencing

293T-B4 attP-containing landing pad cells that were previously described were thawed and passaged twice before recombination. Three replicate wells of a 6 well plate were seeded at 400,000 cells per well and one T25 flask was seeded at 900,000 cells. These were allowed to attach overnight after which they were recombined with the plasmid attB-containing library. All 3 of the 6 wells were transfected with 100 ng of pCAG-NLS-HA-Bxb1 (Addgene #51271) and 1,500 ng of donor attB library whereas the T25 was transfected with 1,000 ng of the Bxb1 containing plasmid and 5,000 ng attB library. Transfection was carried out with lipofectamine 3000 that was constituted in Opti-MEM media. Liposomes were added to cells at ∼30-40% confluence and allowed to incubate overnight. After incubation, media containing the liposomes was removed and fresh media containing puromycin at 0.8 ug/mL was added. After this point puromycin was added to all media in perpetuity to select for successfully recombined cells. After 3 days of puromycin selection, all 4 cultures were trypsinized, pooled, and scaled up to a T75 culture seeding 2.7e6 cells. These were routinely passaged before achieving confluence and selected for a total of 11 days in puromycin before several vials were frozen (5% sterile DMSO v/v stored in the vapor phase of a liquid nitrogen tank) and ∼1 million cells were retained for DNA preparation. Genomic DNA (gDNA) was isolated using the PureLink Genomic DNA Mini kit.

We wanted to validate that the cell library distribution matched the input plasmid library distribution and examine for possible barcode swapping events. We amplified the same barcode containing amplicon off gDNA as we had previously off the plasmid library for NGS. This resulted in 4,649,308 sequences with the same four constant regions described before. Applying the same filtering criteria resulted in 12,269 unique hardcoded – random barcode pairs (Table 2) seen more than once and with more than 95% certainty in the pairing (16,779 in the plasmid library). Of the 8,547 random barcodes seen in both the plasmid library and the gDNA library, only 23 (0.27%) were mapped to different hardcoded barcodes than they previously were when we sequenced off of plasmid DNA. Template switching or otherwise barcode hopping during recombination from the plasmid library into the cell library is therefore exceptionally rare.

### Library and clonal expression validation

In addition to the full library, we performed the same process above to generate a small sub-library and several clonal controls. The clonal controls included EWSR1-FLI1 alone, PAX3-FOXO1 alone, and empty vector control alone. The smaller sub-library included 9 variants pooled: empty vector control, EWSR1-FLI1, EWSR1-ATF1, PAX3-FOXO1, COL1A1-PDGFB, FUS-DDIT3, ETV6-NTRK3, FUS-CREB3L2, and SS18-SSX2. Using these libraries, we validated that the transgenes were induced upon doxycycline expression at the RNA level both in clonal format and in the library format. To do this, 3 biological replicates of each genotype were cultured individually with doxycycline (2 ug/mL) for 48 hours and 3 replicates were cultured with equal volume water. RNA was harvested using Qiagen’s RNeasy Mini kit and the DNAse treatment step was included. One pot qRT-PCR was performed with NEB Luna Universal One-Step RT-qPCR kit and using primers specific to EWSR1-FLI1 or to PAX3-FOXO1. Reactions were monitored using Applied Biosystem’s 7500 Fast Real-Time PCR instrument (Supp Fig 1f) which showed high RNA expression with doxycycline induction for each clonal variant. This corroborated the high protein expression seen by flow cytometry when BFP was used as a proxy prior to transgene recombination (Supp Fig 1a). To determine if expression of any of the sub-library members induced a cell proliferation defect or advantage, we cultured duplicates of the 9-member library in either water or doxycycline for 2 days, 4 days, and 6 days, harvesting gDNA at each time point. We performed the same barcode quantification by sequencing as above and determined that none of the library members altered cell growth (Supp Fig 1h). Taken together, this suggested that a 4 day doxycycline induction of our library would likely be sufficient to induce transgene expression and was unlikely to result in dramatic changes to the library’s distribution.

### Single-cell ATAC and dial-out library preparation

In preparation for sequencing, each library was seeded and allowed to attach over night before being induced with doxycycline (2 ug/mL) for 96 hours. One split was performed at 48 hours. Spear-ATAC is predicated on 10X Genomics single-cell ATAC sequencing. For this work, all sequencing was performed with v2 Chemistry. Nuclei were prepared according to 10X Genomics’ Nuclear Isolation for Single Cell ATAC Sequencing protocol with the following notes. IGEPAL CA-630 was used for the lysis buffer a 10% solution of which was prepared fresh on the day of nuclear isolation using 100% stock (Sigma i8896). After trypsinization and pelleting, cells were washed twice with ice cold PBS + 0.04% BSA before following the remainder of the nuclear isolation protocol. Lysis occurred for 4 minutes which previously was found to be appropriate time for cytoplasmic removal while retaining nuclear membrane architecture by DIC microscopy. Cells were resuspended in diluted nuclei buffer and triplicates were counted before proceeding with Spear-ATAC sequencing as previously published. In all cases, 10,000 nuclei were targeted for capture per 10X chip lane. Spear-ATAC included the following changes to the 10X scATAC library preparation protocol: the 1.2 uL per reaction of 50 uM oSP1735 oligonucleotide is spiked into the GEM generation master mix and the in-GEM PCR was increased from 12 to 15 cycles. Single-cell ATAC library fragment size distribution was analyzed by Tapestation prior to preparing enriched dial-out libraries for perturbation genotyping. The dial-out libraries were again prepared according to the Spear-ATAC protocol with the following changes: linear PCR with biotinylated oSP2053 was increased to 30 total cycles and the exponential PCR with oSP1594 and indexed P7 primers was increased to 18 total cycles. The indices on the P7 primers were increased from 8 bases to 10 bases.

### Single-cell ATAC data processing

All data analysis was carried out on UW Madison’s Center for High Throughput Computing cluster^65^. Sequencing data was converted to fastq files using cellranger-atac mkfastq (10x Genomics, version 2.1.0). Reads were then aligned to the hg38 reference genome and quantified using cellranger-atac count. The current version of Cell Ranger can be accessed here: https://support.10xgenomics.com/single-cell-atac/software/downloads/latest. We used ArchR (version 1.0.0) for most downstream single-cell ATAC-seq analysis (https://greenleaflab.github.io/ArchR_Website/). Fragment files for each sample and the associated cell information file output from CellRanger were passed to ArchR to create Arrow Files. We did not filter doublets because there were few discrete clusters which are necessary for appropriate doublet identification. Next, we filtered for nuclei with TSS enrichment ≥ 7 and number of fragments ≥ 40,000. We did not subset to nuclei with known genotypes prior to generating dimension reduced visualizations reasoning that all nuclei (even if we were unable to genotype them because reads mapping to perturbations are sparse) contained useful information. The UMAP used throughout the manuscript was created with the default parameters though several others were created with various minimum distances and number of neighbors which did not change the interpretation. Most additional analyses—particularly direct differential analysis comparisons between known genotypes—were performed using ArchR.

### Genome annotations

All analysis were performed with the hg38 genome.

### Calling cell perturbation identities with dial-out library sequencing

Dial-out enrichment libraries with which to call perturbation identities in individual nuclei were sequenced as in Spear-ATAC. However, in this case random barcodes were sequenced (instead of gRNAs) and those were then mapped back to hardcoded barcodes and therefore the encoded protein-coding variant. Random barcodes were identified by grepping for several constant sequences to ensure that amplicons of the correct format were analyzed: “AAATCCAAGC”, “CCAGAGCATG”, “CAAGGTGGTT”, and “ATACTGATTC”. Random barcodes were then isolated alongside their corresponding cell barcodes from the 10X library construction process (CBCs). We filtered on 10X CBCs that were seen at least 3 times and then only retained pairs of 10X CBC – Random barcode for which ≥ 90% of the reads were the same pair (i.e. a 10X CBC that was associated with equipoise to multiple random barcodes was not retained). Random barcodes from the dial-out library were then matched to random barcodes seen from sequencing the library from gDNA previously. For those random barcodes that matched one of the stringent set of random barcodes previously defined, we were able to confidently identify the protein-coding variant present for the associated cell barcode. The remainder of cell barcodes that were unidentified were not used in downstream analysis.

### Comparing single-cell pseudobulk data to external references

We sought to determine how differential peaks in cells containing variants from our library compared to significant peaks from known human cell types. To do this, we created BED files for peaks increased in accessibility for each fusion compared to empty vector control (FDR < 0.1 and log_2_(fold change) ≥ 1) using ArchR’s ability to directly compare pseudobulk replicates of two groups. These BED files were then passed to ChIP-Atlas’s Enrichment Analysis feature with settings: Experiment type = ATAC-Seq, Cell type class = All cell types, Threshold for significance = 200, and dataset B (control dataset) = random permutations of dataset A 100 times. We combined the resulting data frames and for the purpose of visualization we subsetted to those overlaps which had -log(p value) > 3 and for which the fraction of fusion peaks overlapping with the given known cell type was above 0.2 as these are the likely most meaningful overlaps.

### Down-sampling single-cell ATAC data

To ascertain the effect of capturing fewer cells on the resulting data, we systematically down-sampled the number of nuclei captured for several fusions and controls. In these cases, we created random samples of the existing genotyped cells ranging from 10 nuclei to the total number of nuclei captured for that genotype. In the case of empty vector control, this was performed at intervals of 100 nuclei whereas for all fusions it was performed at intervals of 20 nuclei. For each random sample, we used ArchR to directly compare the down-sampled datasets for each variant to empty vector control. We then determined the Pearson correlation between the log_2_(fold change) for each peak in the down-sampled comparison to that of the original comparison. Similarly, we determined the total number of differentially accessible peaks in the down-sampled comparison as a fraction of the total number of differentially accessibile peaks in the original comparison with all nuclei.

## Data availability

All sequencing data was deposited in the Gene Expression Omnibus (GEO) which will become publicly available at the time of publication. Plasmids generated in this study are available from the lead contact (S.R.) without restriction. Any other relevant data are available from the authors upon reasonable request.

## Code availability

We are hosting a GitHub website that includes the main code used in this study (mfrenkel16/OncofusionPRODATAC/). Any other code is available upon request.

## Author contributions

M.F. conceived of the study, performed all experiments, analyzed and interpreted all data, and wrote the original manuscript. M.L.A.H. mined fusion databases to create the *in silico* library and edited the manuscript. Z.M. supervised the work, provided helpful biological insight, acquired funding, and edited the manuscript. S.R. supervised the work, provided valuable guidance throughout the study, acquired funding, and edited the manuscript. All authors read and approved of the final manuscript.

## Acknowledgments

M.F. was supported by an NIH T32 fellowship (T32HG2760-17). This work was partially supported by the NIH Director’s New Innovator Award DP2GM132682 (S.R.) and UWCCC Core grant P30CA014520 (Z.M.). We would also like to thank Rein in Sarcoma for a charitable donation that partially supported this work. We thank both Kenneth Matreyek and Peter Lewis for sharing plasmids and cells. We thank Silas Miller for his helpful discussion regarding barcoding. Most DNA amplicon sequencing was performed at the University of Wisconsin – Madison Biotechnology Center’s DNA Sequencing Facility (Research Resource Identifier – RRID:SCR_017759) as was all Tapestation analysis of single-cell library fragment distributions. Microscopy was performed at the University of Wisconsin-Madison Biochemistry Optical Core which was established with support from the University of Wisconsin-Madison Department of Biochemistry Endowment. Fluorescence activated cell sorting and flow cytometry were performed at UW-Madison Flow Cytometry core.

## Ethics declaration

The authors declare no competing interests.

## References

1. Przybyla, L. & Gilbert, L. A. A new era in functional genomics screens. Nat. Rev. Genet. 1–15 (2021) doi:10.1038/s41576-021-00409-w.

2. Findlay, G. M. Linking genome variants to disease: scalable approaches to test the functional impact of human mutations. Hum. Mol. Genet. 30, R187–R197 (2021).

3. Findlay, G. M. et al. Accurate classification of BRCA1 variants with saturation genome editing. Nature 562, 217–222 (2018).

4. Tewhey, R. et al. Direct Identification of Hundreds of Expression-Modulating Variants using a Multiplexed Reporter Assay. Cell 165, 1519–1529 (2016).

5. van Arensbergen, J. et al. High-throughput identification of human SNPs affecting regulatory element activity. Nat. Genet. 51, 1160–1169 (2019).

6. Doench, J. G. Am I ready for CRISPR? A user’s guide to genetic screens. Nat. Rev. Genet. 19, 67–80 (2018).

7. Fowler, D. M. & Fields, S. Deep mutational scanning: a new style of protein science. Nat. Methods 11, 801–807 (2014).

8. Replogle, J. M. et al. Combinatorial single-cell CRISPR screens by direct guide RNA capture and targeted sequencing. Nat. Biotechnol. 1–8 (2020) doi:10.1038/s41587-020-0470-y.

9. Adamson, B. et al. A Multiplexed Single-Cell CRISPR Screening Platform Enables Systematic Dissection of the Unfolded Protein Response. Cell 167, 1867–1882.e21 (2016).

10. Dixit, A. et al. Perturb-Seq: Dissecting Molecular Circuits with Scalable Single-Cell RNA Profiling of Pooled Genetic Screens. Cell 167, 1853–1866.e17 (2016).

11. Datlinger, P. et al. Pooled CRISPR screening with single-cell transcriptome readout. Nat. Methods 14, 297–301 (2017).

12. Kim, H. S. et al. Direct measurement of engineered cancer mutations and their transcriptional phenotypes in single cells. Nat. Biotechnol. 1–9 (2023) doi:10.1038/s41587-023-01949-8.

13. Ursu, O. et al. Massively parallel phenotyping of coding variants in cancer with Perturb-seq. Nat. Biotechnol. 40, 896–905 (2022).

14. Pierce, S. E., Granja, J. M. & Greenleaf, W. J. High-throughput single-cell chromatin accessibility CRISPR screens enable unbiased identification of regulatory networks in cancer. Nat. Commun. 12, 2969 (2021).

15. Mertens, F., Johansson, B., Fioretos, T. & Mitelman, F. The emerging complexity of gene fusions in cancer. Nat. Rev. Cancer 15, 371–381 (2015).

16. F, M., Cr, A. & F, M. Gene fusions in soft tissue tumors: Recurrent and overlapping pathogenetic themes. Genes. Chromosomes Cancer 55, (2016).

17. Gryder, B. E. et al. PAX3–FOXO1 Establishes Myogenic Super Enhancers and Confers BET Bromodomain Vulnerability. Cancer Discov. 7, 884–899 (2017).

18. Riggi, N. et al. EWS-FLI1 utilizes divergent chromatin remodeling mechanisms to directly activate or repress enhancer elements in Ewing sarcoma. Cancer Cell 26, 668–681 (2014).

19. Boulay, G. et al. Cancer-Specific Retargeting of BAF Complexes by a Prion-like Domain. Cell 171, 163–178.e19 (2017).

20. Jang, Y. E. et al. ChimerDB 4.0: an updated and expanded database of fusion genes. Nucleic Acids Res. 48, D817–D824 (2020).

21. Sweeney, N. P. & Vink, C. A. The impact of lentiviral vector genome size and producer cell genomic to gag-pol mRNA ratios on packaging efficiency and titre. Mol. Ther. Methods Clin. Dev. 21, 574–584 (2021).

22. Milone, M. C. & O’Doherty, U. Clinical use of lentiviral vectors. Leukemia 32, 1529–1541 (2018).

23. Xie, S., Cooley, A., Armendariz, D., Zhou, P. & Hon, G. C. Frequent sgRNA-barcode recombination in single-cell perturbation assays. PLOS ONE 13, e0198635 (2018).

24. Adamson, B., Norman, T. M., Jost, M. & Weissman, J. S. Approaches to maximize sgRNA-barcode coupling in Perturb-seq screens. 298349 https://www.biorxiv.org/content/10.1101/298349v1 (2018) doi:10.1101/298349.

25. Feldman, D., Singh, A., Garrity, A. J. & Blainey, P. C. Lentiviral co-packaging mitigates the effects of intermolecular recombination and multiple integrations in pooled genetic screens. 262121 Preprint at 10.1101/262121 (2018).

26. Parekh, U. et al. Mapping Cellular Reprogramming via Pooled Overexpression Screens with Paired Fitness and Single-Cell RNA-Sequencing Readout. Cell Syst. 7, 548–555.e8 (2018).

27. Hill, A. J. et al. On the design of CRISPR-based single-cell molecular screens. Nat. Methods 15, 271–274 (2018).

28. Matreyek, K. A., Stephany, J. J., Chiasson, M. A., Hasle, N. & Fowler, D. M. An improved platform for functional assessment of large protein libraries in mammalian cells. Nucleic Acids Res. 48, e1 (2020).

29. Ursu, O. et al. Massively parallel phenotyping of variant impact in cancer with Perturb-seq reveals a shift in the spectrum of cell states induced by somatic mutations. bioRxiv 2020.11.16.383307 (2020) doi:10.1101/2020.11.16.383307.

30. Sánchez-Molina, S. et al. RING1B recruits EWSR1-FLI1 and cooperates in the remodeling of chromatin necessary for Ewing sarcoma tumorigenesis. Sci. Adv. 6, eaba3058 (2020).

31. Deng, Q. et al. Oncofusion-driven de novo enhancer assembly promotes malignancy in Ewing sarcoma via aberrant expression of the stereociliary protein LOXHD1. Cell Rep. 39, 110971 (2022).

32. Manceau, L. et al. Divergent transcriptional and transforming properties of PAX3-FOXO1 and PAX7-FOXO1 paralogs. PLOS Genet. 18, e1009782 (2022).

33. Orth, M. F. et al. Systematic multi-omics cell line profiling uncovers principles of Ewing sarcoma fusion oncogene-mediated gene regulation. Cell Rep. 41, 111761 (2022).

34. Tate, J. G. et al. COSMIC: the Catalogue Of Somatic Mutations In Cancer. Nucleic Acids Res. 47, D941–D947 (2019).

35. Granja, J. M. et al. ArchR is a scalable software package for integrative single-cell chromatin accessibility analysis. Nat. Genet. 53, 403–411 (2021).

36. Jost, M. et al. Titrating gene expression using libraries of systematically attenuated CRISPR guide RNAs. Nat. Biotechnol. 38, 355–364 (2020).

37. Hao, Y. et al. Integrated analysis of multimodal single-cell data. Cell 184, 3573–3587.e29 (2021).

38. Zhang, Y. et al. Model-based Analysis of ChIP-Seq (MACS). Genome Biol. 9, R137 (2008).

39. Sunkel, B. D. et al. Evidence of pioneer factor activity of an oncogenic fusion transcription factor. iScience 24, 102867 (2021).

40. AACR Project GENIE Consortium. AACR Project GENIE: Powering Precision Medicine through an International Consortium. Cancer Discov. 7, 818–831 (2017).

41. Chang, W.-I. et al. Molecular Targets for Novel Therapeutics in Pediatric Fusion-Positive Non-CNS Solid Tumors. Front. Pharmacol. 12, (2022).

42. Perry, J. A., Seong, B. K. A. & Stegmaier, K. Biology and Therapy of Dominant Fusion Oncoproteins Involving Transcription Factor and Chromatin Regulators in Sarcomas. Annu. Rev. Cancer Biol. 3, 299–321 (2019).

43. Möller, E. et al. EWSR1-ATF1 dependent 3D connectivity regulates oncogenic and differentiation programs in Clear Cell Sarcoma. Nat. Commun. 13, 2267 (2022).

44. Johnson, K. M. et al. Role for the EWS domain of EWS/FLI in binding GGAA-microsatellites required for Ewing sarcoma anchorage independent growth. Proc. Natl. Acad. Sci. U. S. A. 114, 9870–9875 (2017).

45. Johnson, K. M., Taslim, C., Saund, R. S. & Lessnick, S. L. Identification of two types of GGAA-microsatellites and their roles in EWS/FLI binding and gene regulation in Ewing sarcoma. PLOS ONE 12, e0186275 (2017).

46. Guillon, N. et al. The oncogenic EWS-FLI1 protein binds in vivo GGAA microsatellite sequences with potential transcriptional activation function. PloS One 4, e4932 (2009).

47. Li, Z. et al. ETV6-NTRK3 fusion oncogene initiates breast cancer from committed mammary progenitors via activation of AP1 complex. Cancer Cell 12, 542–558 (2007).

48. Przybyl, J. et al. Gene expression profiling of low-grade endometrial stromal sarcoma indicates fusion protein-mediated activation of the Wnt signaling pathway. Gynecol. Oncol. 149, 388–393 (2018).

49. Gordon, A. T. et al. A novel and consistent amplicon at 13q31 associated with alveolar rhabdomyosarcoma. Genes. Chromosomes Cancer 28, 220–226 (2000).

50. Yoon, J. W., Lamm, M., Chandler, C., Iannaccone, P. & Walterhouse, D. Up-regulation of GLI1 in vincristine-resistant rhabdomyosarcoma and Ewing sarcoma. BMC Cancer 20, 511 (2020).

51. Birdsey, G. M. et al. The endothelial transcription factor ERG promotes vascular stability and growth through Wnt/β-catenin signaling. Dev. Cell 32, 82–96 (2015).

52. Brcic, I. et al. Implementation of Copy Number Variations-Based Diagnostics in Morphologically Challenging EWSR1/FUS::NFATC2 Neoplasms of the Bone and Soft Tissue. Int. J. Mol. Sci. 23, 16196 (2022).

53. Deplus, R. et al. TMPRSS2-ERG fusion promotes prostate cancer metastases in bone. Oncotarget 8, 11827–11840 (2016).

54. Parviz, F. et al. Hepatocyte nuclear factor 4alpha controls the development of a hepatic epithelium and liver morphogenesis. Nat. Genet. 34, 292–296 (2003).

55. Zhang, B. et al. Proteogenomic characterization of human colon and rectal cancer. Nature 513, 382–387 (2014).

56. Weinstein, J. N. et al. The Cancer Genome Atlas Pan-Cancer analysis project. Nat. Genet. 45, 1113–1120 (2013).

57. Davis, R. B., Kaur, T., Moosa, M. M. & Banerjee, P. R. FUS oncofusion protein condensates recruit mSWI/SNF chromatin remodeler via heterotypic interactions between prion-like domains. Protein Sci. Publ. Protein Soc. 30, 1454–1466 (2021).

58. Backman, J. D. et al. Exome sequencing and analysis of 454,787 UK Biobank participants. Nature 1–10 (2021) doi:10.1038/s41586-021-04103-z.

59. Lancaster, A. K., Nutter-Upham, A., Lindquist, S. & King, O. D. PLAAC: a web and command-line application to identify proteins with prion-like amino acid composition. Bioinformatics 30, 2501–2502 (2014).

60. Farag, M., Borcherds, W. M., Bremer, A., Mittag, T. & Pappu, R. V. Phase Separation in Mixtures of Prion-Like Low Complexity Domains is Driven by the Interplay of Homotypic and Heterotypic Interactions. 2023.03.15.532828 Preprint at 10.1101/2023.03.15.532828 (2023).

61. Boncella, A. E. et al. Composition-based prediction and rational manipulation of prion-like domain recruitment to stress granules. Proc. Natl. Acad. Sci. 117, 5826–5835 (2020).

62. Sprunger, M. L. & Jackrel, M. E. Prion-Like Proteins in Phase Separation and Their Link to Disease. Biomolecules 11, 1014 (2021).

63. Wang, Y. et al. Dissolution of oncofusion transcription factor condensates for cancer therapy. Nat. Chem. Biol. (2023) doi:10.1038/s41589-023-01376-5.

64. Frankish, A. et al. GENCODE 2021. Nucleic Acids Res. 49, D916–D923 (2021).

65. Center for High Throughput Computing. Center for High Throughput Computing. (2006) doi:10.21231/GNT1-HW21.

