## Supplemental figures for "Discovering chromatin dysregulation induced by protein-coding perturbations at scale"

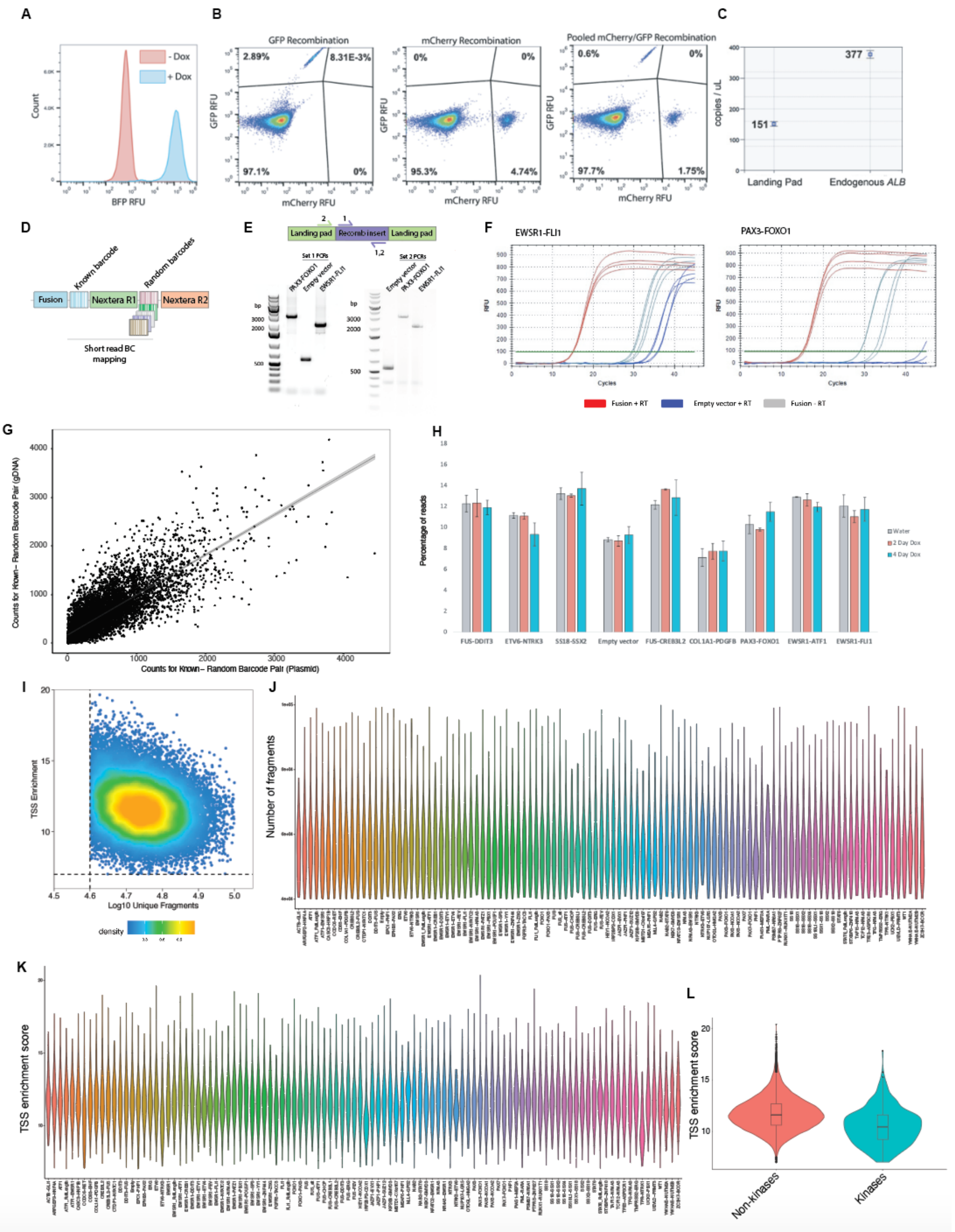
A
- Dox
+ Dox
Count
BFP RFU
B
GFP Recombination
2.89%
8.31E-3%
97.1%
0%
mCherry Recombination
0%
0%
95.3%
4.74%
Pooled mCherry/GFP Recombination
0.6%
0%
97.7%
1.75%
GFP RFU
mCherry RFU
C
377
151
copies / uL
Landing Pad
Endogenous ALB
D
Fusion
Known barcode
Random barcodes
Nextera R1
Nextera R2
Short read BC mapping
E
Landing pad
Recomb insert
Landing pad
Set 1 PCRs
Set 2 PCRs
PAX3-FOXO1
Empty vector
EWSR1-FLI1
bp
3000
2000
500
F
EWSR1-FLI1
PAX3-FOXO1
RFU
Cycles
Fusion + RT
Empty vector + RT
Fusion - RT
G
Counts for Known– Random Barcode Pair (gDNA)
Counts for Known– Random Barcode Pair (Plasmid)
H
Percentage of reads
Water
2 Day Dox
4 Day Dox
FUS-DDIT3
ETV6-NTRK3
SS18-SSX2
Empty vector
FUS-CREB3L2
COL1A1-PDGFB
PAX3-FOXO1
EWSR1-ATF1
EWSR1-FLI1
I
TSS Enrichment
Log10 Unique Fragments
density
J
Number of fragments
K
TSS enrichment score
L
TSS enrichment score
Non-kinases
Kinases

**Supplemental Figure 1: A generalizable platform for screening protein-coding variants including oncofusions.** (A): Histogram of flow cytometry of clonal landing pad 293T cells with and without doxycycline based on blue fluorescence protein (BFP) expression. (B) Test recombination of either pure green fluorescence protein (GFP) containing donor plasmid, pure mCherry containing plasmid, or pooled mCherry and GFP plasmid in equimolar ratio. (C) Droplet digital PCR off of genomic DNA of clonal landing pad 293T cells. One primer set was specific for the landing pad site and the other for the endogenous *ALB* gene. (D) Barcoding scheme for identifying variants in nuclei. Fusions are synthesized with known (hardcoded) barcodes followed by golden gate which appends a random barcode flanked by Nextera adapters. The random barcode and known barcode can be sequenced with one short-read NGS run. (E) PCR validation of clonal oncofusion and control recombination into 293T landing pad cells. (F) Clonal validation of doxycycline-induced expression of oncofusions compared to control by qRT-PCR. (G) Correlation between distribution of random barcodes after inserting into genomic DNA (gDNA) and in plasmid DNA library (H) Small scale test of oncofusion distribution over time in a pooled library format. (I) TSS enrichment score versus number of unique fragments sequenced for all nuclei captured with Spear-ATAC after filtering. (J) Distribution of number of fragments captured for each nuclei associated with each genotype. (K) Distribution of TSS enrichment score for each nuclei associated with each genotype. (L) Distribution of TSS enrichment score separated by whether the fusion contains a kinase or not.

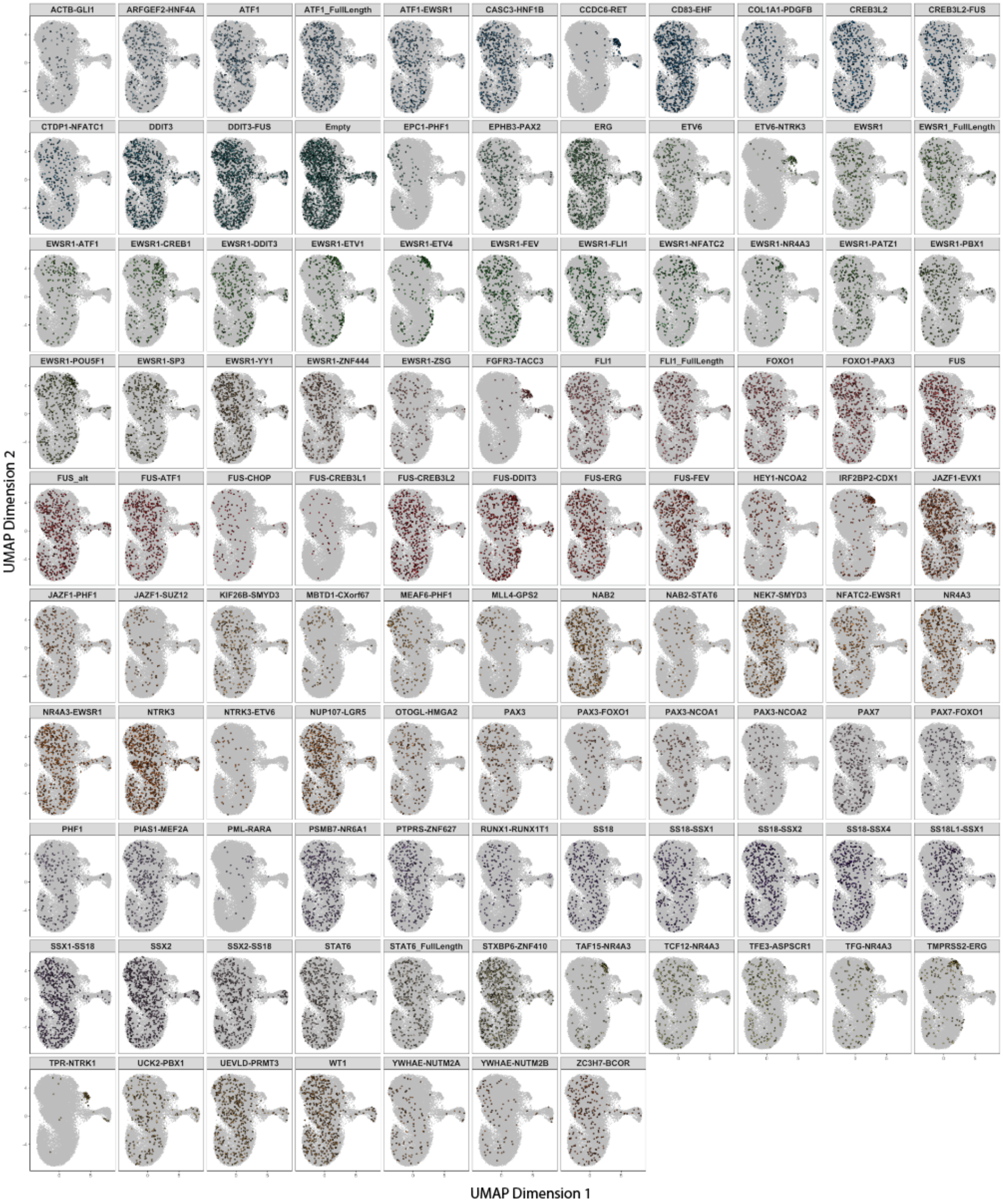


**Supplemental Figure 2: Cell-state distributions in UMAP space for each variant.** All 35,086 nuclei are displaced in UMAP space after latent semantic indexing. Within each panel, nuclei containing a given genotype are highlighted.

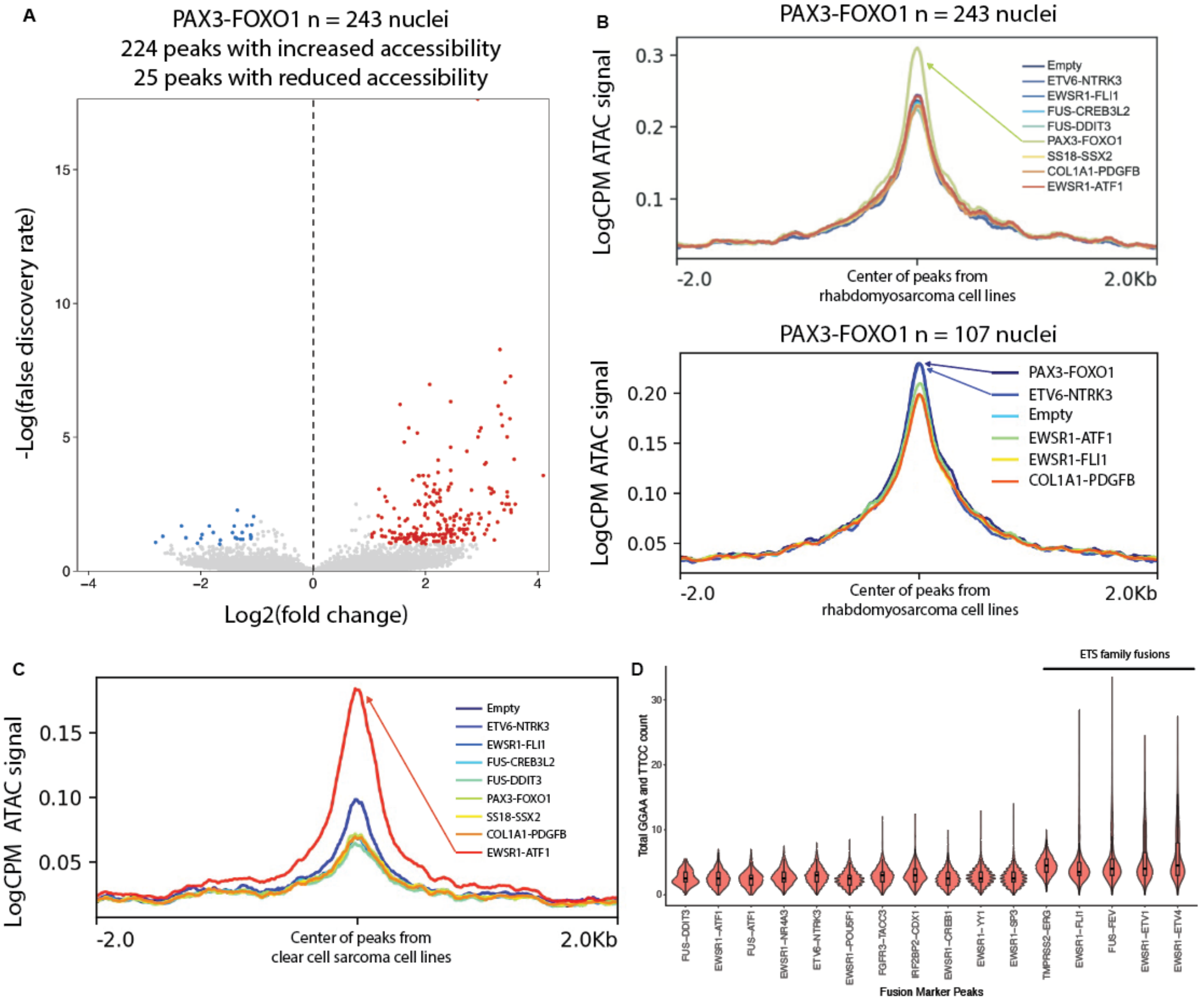


**Supplemental Figure 3: Oncofusion overexpression in 293T recapitulates known cancer-specific and biochemical signatures.** (A) Volcano plot of -Log(false discovery rate) versus Log2(fold change) comparing PAX3-FOXO1 expressing cells to empty vector control at all peaks from the smaller scale experiment. (B) LogCPM normalized signal after aggregating all cells expressing PAX3-FOXO1 or controls. Center of peaks is defined by loci that are PAX3-FOXO1 bound in rhabdomyosarcoma cells from Sunkel et al. *iScience.* **2021.** Top is data from a small scale experiment with only 9 total variants and 243 nuclei capture representing PAX3-FOXO1 whereas the bottom is from the large scale experiment with 112 variants and 107 nuclei representing PAX3-FOXO1. (C) LogCPM normalized signal after aggregating cells based on genotype. Center of peaks is defined by loci that are EWSR1-ATF1 bound in clear cell sarcoma cell lines from Möller et al. *Nat Commun.* **2022.** (D) Total GGAA and TTAA count in differentially accessible peaks for all ETS family fusions and several controls.

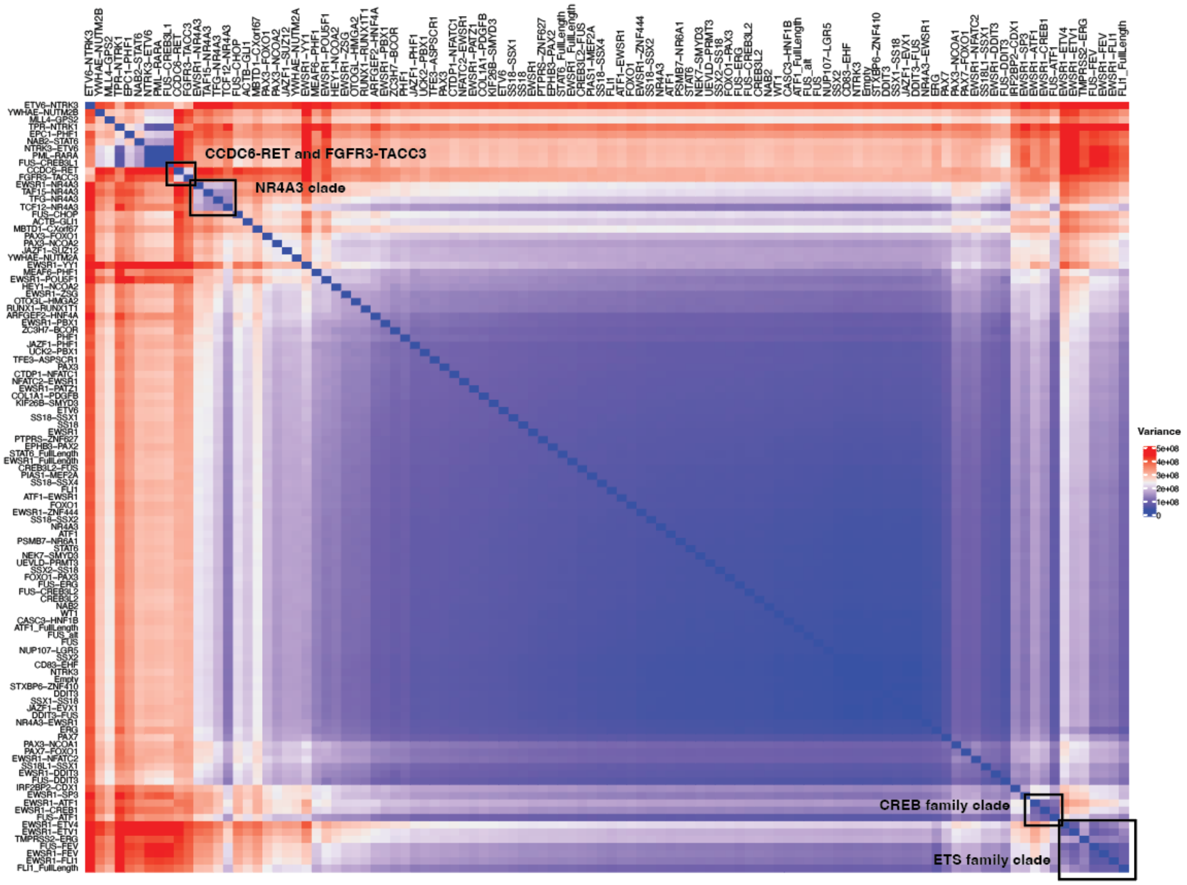


**Supplement Figure 4: Robustness of oncofusion classification.** Distances for all pairwise comparisons between variants in the library were calculated with 4 distance metrics (Pearson, Spearman, Euclidean, and Manhattan). The variance among these measurements is plotted above and clustered.

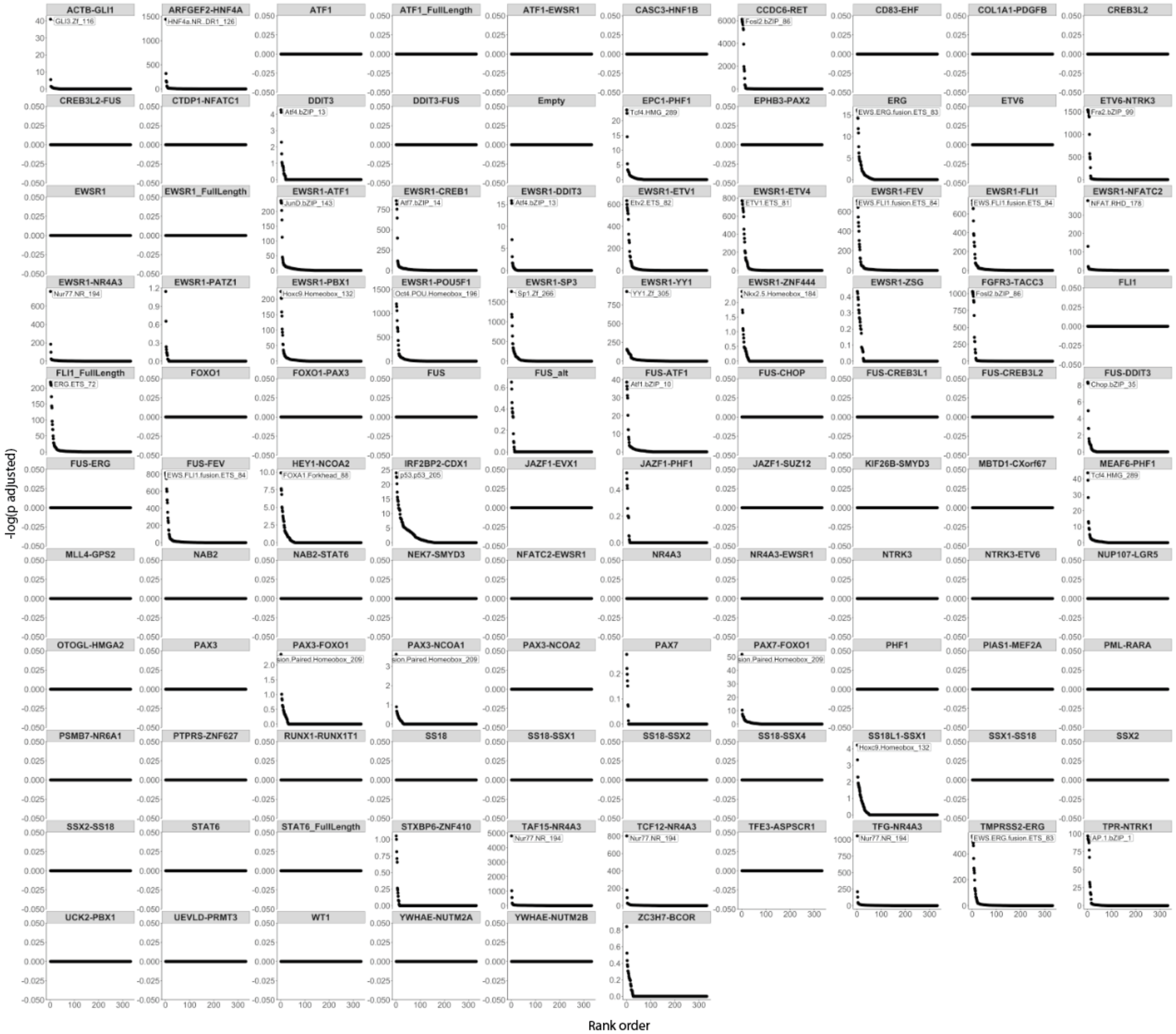


**Supplement Figure 5: Recapitulation of known oncofusion biology.** DNA motif enrichment was calculated for peaks with increased differential accessibility when comparing each variant in the library to empty vector control.

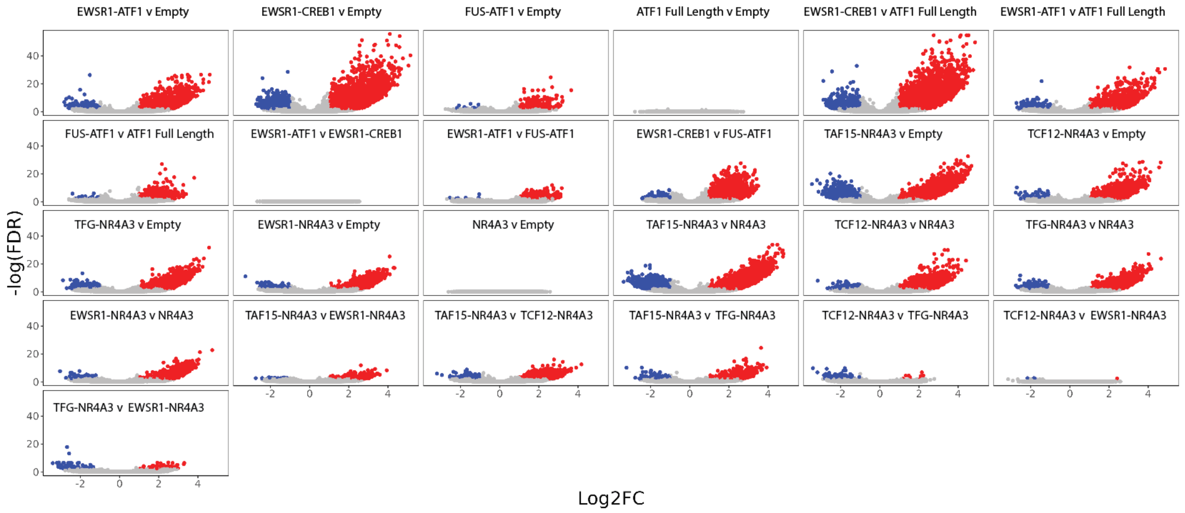


**Supplement Figure 6: Many oncofusions exhibit gain of function chromatin pioneering.** Volcano plot comparisons among oncofusion proteins as well as between oncofusions and domain controls or empty vector control in pseudobulk. Red peaks are significantly increased in accessibility and blue peaks are significantly reduced in accessibility.
